# Common but different: An ERP study of single- and multi-source interference processing in MSIT

**DOI:** 10.1101/2025.03.05.641477

**Authors:** Katarzyna Paluch, Ingrida Antonova, Jan Nikadon, Patrycja Dzianok, Katarzyna Jurewicz, Jakub Wojciechowski, Ewa Kublik

## Abstract

Flexible behavior often requires processing of complex, interfering information. Research has investigated conflict-related brain processes mostly using single tasks which hindered direct comparison of different interference types. Thus, the question if they are resolved by a common mechanism or by a set of different, task-specific mechanisms remains open. In this study, we used event-related potentials (ERPs) to examine the spatio-temporal dynamics of cognitive control across Simon, flanker, multi-source and no-conflict conditions. Our findings reveal that all trial types engaged the same sequence of processing stages, as indicated by common ERP waveforms and consistent number and order of microstates across conditions. However, the intensity and duration of these common stages scaled with difficulty of the conflict task (as measured by RTs and accuracy) from Simon to flanker to multi-source interference. Flanker conflict uniquely influenced early ERP components strongly engaging the dorsal attentional system and visual areas, likely due to demands posed by the presence of flanker distractors. Later ERP components (with sources including ventral attention and somatomotor network areas) were affected by both conflicts. Accordingly, when flanker and Simon conflicts were presented together, early processes lineary summed up, but there was an interaction at the later stage of processing paralleling nonlinear drop of accuracy in a multi-conflict condition. Our study provides novel insights into the neural dynamics underlying cognitive control engaged across different conflict types and their interaction. The use of source analysis allowed us to ground ERP-based findings in the wider context of studies, including those using neuroimaging techniques.

**Highlights:**

- The same processing stages, in same order are evoked in control and conflict trials
- Their intensity/duration scale with conflict behavioral difficulty
- Flanker conflict demands enhanced early attentional (DAN) and visual processing
- Flanker and Simon interact at the late-stage processing involving VAN and SM networks
- Flanker and Simon interaction is sublinear

## 1. Introduction

The ability to select relevant and inhibit irrelevant information is necessary for complex, adaptive behavior. The process is possible thanks to recruitment of the brain’s cognitive control mechanisms. Decades of research have led to the development of numerous paradigms aiming to study cognitive control, which are jointly referred to as conflict tasks. Among the most popular ones are the Stroop task (Stroop, 1935), the Simon task (Simon & Rudell, 1967) and the Eriksen flanker task (Eriksen & Eriksen, 1974). These tasks share a common structure in which participants are required to respond to a particular feature of a stimulus, while ignoring competing, interfering features and their associated responses. In the Simon task, stimulus position on the screen is incongruent with the required response (e.g. a stimulus presented on the left side of the screen requires a response with the right hand). In the Eriksen flanker task, subjects respond to a target stimulus, which is surrounded by salient distractors (flankers) associated with different responses. These two interference types were combined in the Multi-Source Interference Task (MSIT, Bush & Shin, 2006) which was designed to maximize the demand for cognitive control engagement.

Theoretically, various conflict-inducing tasks could be resolved either by a common control mechanism or by multiple, task-specific mechanisms. The literature offers conflicting (nomen omen) conclusions on this topic. Some functional magnetic resonance imaging (fMRI) studies supported the hypothesis of general conflict resolution mechanism (Botvinick et al., 2004; Peterson et al., 2002). Others suggest task-specific activations (Egner et al., 2007; Kim et al., 2010) or a combination of both common and task specific processes (Li et al., 2017; Wager et al., 2005; Wojciechowski et al., 2024). Electroencephalography (EEG) studies, on the other hand, report that different event-related potential (ERP) components are modulated by different types of conflict (for review see: (Larson et al., 2014; Pires et al., 2014). Studies using the Eriksen flanker task (Folstein & Van Petten, 2008; van Veen & Carter, 2002) most often report N2 component, a negative, fronto-central potential peaking around 200-300 ms after stimulus onset. Tasks with semantic components, such as the Stroop task, typically elicit the N400/N450 component. This component shares a fronto-central topography with the N2 but occurs later after stimulus onset (Chuderski et al., 2016; West, 2003). Notably, the N400/N450 is absent in non-linguistic tasks, such as the Simon or Eriksen flanker tasks, where conflict-related effects are instead observed in the P3 component (Heidlmayr et al., 2020). The final conflict-related ERP is the Slow/Sustained Potential (SP), which occurs around 500-800 ms after stimulus onset. This potential is characterized by frontal negativity and parietal positivity and has been reported in a variety of conflict tasks, including the Stroop (West, 2003; West et al., 2005), the flanker (Donohue et al., 2016), and the Simon (Chen & Melara, 2009) tasks.

The more complex picture arising from EEG compared to the fMRI studies may partially result from the different time resolution of the two methods, with EEG being able to discriminate the consecutive stages of cognitive processes and detect subtle differences in their onsets and durations. On the other hand, higher sensitivity may result in a ‘noisier’ picture with EEG picking up task-specific factors unrelated to interference but determining which ERP components are affected when a particular task is used. To resolve this issue, a direct comparison of ERPs across different tasks is required. Unfortunately, almost all existing studies rely on a single task, which restricts comparisons to those between studies and prevents conclusive interpretation of differences in brain activity.

Few ERP studies have directly compared different types of interference within a single study using the same stimuli (Frühholz et al., 2011a; Korsch et al., 2016; Rey-Mermet et al., 2019a; Scrivano & Kieffaber, 2022; Xie et al., 2020) or two sets of stimuli presented in separate blocks but within a single session (Donohue et al., 2016). Studies combining flanker and Simon interference (Frühholz et al., 2011a; Korsch et al., 2016; Scrivano & Kieffaber, 2022) showed flanker-specific effects during early processing stages (N2/P2). However, findings for later stages are inconsistent, with some studies reporting Simon-specific effects on P3 (Frühholz et al., 2011a) and others showing that both Simon and flanker conflicts impacted the P3 component (Korsch et al., 2016; Scrivano & Kieffaber, 2022). Similar observations were made in studies combining flanker and Stroop conflicts in which flanker-specific effects were observed during early processing stages (P2; Donohue et al., 2016; Rey-Mermet et al., 2019a). However, conclusions about the later processing stages differed, with Stroop-specific effects reported for an N450-like component (Rey-Mermet et al., 2019a; Scrivano & Kieffaber, 2022) the N450-like component but with task-specific timing (Donohue et al., 2016). The inconsistencies in existing reports may be partially due to methodological differences in: channel focus, time windows, reporting of difference-waves rather than raw potentials (Scrivano & Kieffaber, 2022) and variation in components definitions, where the same labels are applied to waves of different latencies. To facilitate synthesis, studies with comprehensive reports of results across all electrodes and entire trial durations are needed.

Another unresolved question is what happens when multiple conflicts are present simultaneously (a situation likely to arise in everyday life). Are they resolved independently (resulting in linear summation of time or activity area/power), or do the processes involved in their resolution overlap (leading to interactions)? Paradigms using single and combined interference types within the same experiment (i.e. above mentioned reports) have shown nonlinear effects in behavioral results but no interaction effects/nonlinearities in ERP measures (Frühholz et al., 2011b; Rey-Mermet et al., 2019b). This discrepancy may be attributed to differences in the analysis of behavioral and ERP data. Behavioral data were typically analyzed using two-way ANOVA, where both conflicts were treated as separate factors. However, this approach has rarely been applied to ERP data (e.g. Rey-Mermet et al., 2019a). Previous studies (Korsch et al., 2016) often employed multivariate analyses approach, involving multiple factors unrelated to conflict type (e.g. electrodes positions, response time, subjects age) which may have reduced their sensitivity to smaller effects of different conflicts interactions. On the other hand, the studies, which focused on multiple conflict interactions and directly compared the multi-conflict effects in additive model *versus* real data from MSIT task reported nonlinearities at both behavioral and neuronal level (Arif et al., 2023; Wiesman & Wilson, 2020). However, they focused on the oscillatory activity and did not report ERFs (event-related fields, MEG equivalent of ERPs) results.

We aimed to address these questions by employing an extended variant of the Multi-Source Interference Task (Sheth et al., 2012) with four types of trials: (1) no-conflict, (2) Simon-only conflict, (3) flanker-only conflict and (4) multi-source interference trials with concurrent Simon and flanker conflicts. Until now, the basic variant of MSIT (i.e. without single-conflict conditions) was sporadically used in EEG studies, with reports limited to individual preselected channels and time windows (González-Villar & Carrillo-de-la-Peña, 2017; Robertson et al., 2014; Widge et al., 2019). The extended variant of MSIT (i.e. with single-conflict conditions) has only been used in MEG studies (Arif et al., 2023; Wiesman et al., 2020; Wiesman & Wilson, 2020). Thus, the presented work is the first to report a complete set of ERPs and their brain sources evoked by extended MSIT.

We used cluster based permutation approach to verify task-related amplitude differences in ERPs waves on all electrodes through the entire trial duration, and we applied the microstate analysis (Lehmann et al., 1987) to evaluate the temporal dynamics and intensity of semi-stationary patterns of brain activity (see review by Schiller et al., 2024). Microstate analysis has previously indicated engagement of the same processes in congruent and incongruent conditions for IAT (Schiller et al., 2016) and Stroop tasks (Ruggeri et al., 2019). However, it is not clear if common or conflict-type specific processes are involved during resolution of other interference types, e.g. flanker and Simon. New findings regarding sequence and dynamics of microstates activation are even more important to clarify conflicting reports (e.g. whether it is the same processing stage despite various ERP names or not) and tackle a question about sequential/simultaneous processing and mental states involved. Finally, to enhance synthesis of fMRI and EEG findings, which are largely developed in parallel, we localized brain sources for each of the identified microstates and referred obtained source activity maps to the functional large scale brain networks (Yeo et al., 2011) to facilitate their functional interpretation. As a final analytical step, we tested whether differences in the intensity and duration of earlier microstates contribute to the differences in the intensity and duration of later ones and correlate with interpersonal differences in task performance.

## 2. Materials and Methods

### 2.1 Participants

41 healthy adults (22 females, 19 males, mean age 24.63, SD = 4.17 years) participated in the study. Inclusion criteria were right-handedness, age from 20 to 35 years, normal or corrected to normal vision, medication-free, and no history of head trauma, neurological and psychiatric conditions, chronic disease. The EEG session was a part of the larger project which included fMRI and psychological testing sessions (Wojciechowski et al., 2024). Thus, exclusion criteria were adjusted to fMRI safety requirements and included pregnancy, wearing braces, having metal implants or heart stimulators. In addition to self-report, right-handedness of all participants was confirmed by a short version of Edinburgh Handedness Inventory (Oldfield, 1971). The experiment was approved by the local ethics committee and was conducted in accordance with statutes of the Declaration of Helsinki. All participants provided written informed consent for participation in the study.

### 2.2 Stimuli and task

We adapted the Multi-Source Interference Task (MSIT; (Bush et al., 2003; Bush & Shin, 2006) with the modification introduced by (Sheth et al., 2012). The modified MSIT consisted of four conditions each with a particular type of cognitive conflict involved: 1) control/no-interference, 2) Simon, 3) Flanker, and 4) multi-source (combined Flanker and Simon) conflict (see Fig. 1 for example stimuli). Participants were instructed to identify the target digit by pressing a response button corresponding to the value of the target digit and ignore its spatial position. Responses were provided using keys “1”, “2” and “3” located on the numerical keyboard and participants had to press them with the index, middle and ring finger of the right hand, respectively. We instructed subjects to provide answers as quickly as possible, but not to sacrifice correctness and to avoid a second response after they made a mistake.

**Figure 1.**
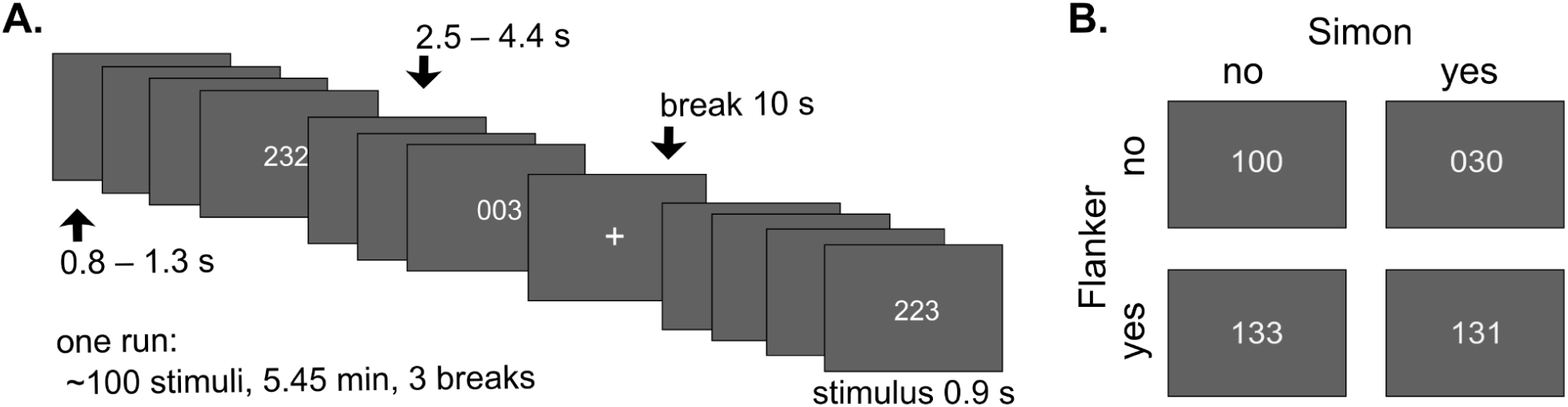
Experimental procedure and example stimuli. A) Single trial consisted of a 900 ms stimulus presentation followed by an inter-stimulus interval (800-1300 ms or 2500-4400 ms, the longer version after each miniblock of 3-4 trials). Short break (indicated by a white cross) was introduced after 9-10 mini-blocks and a longer rest period was allowed after a run of ∼100 trials. The duration of a rest was not defined and ended by a participant executing a key-press. The full experiment consisted of four runs. B) Example stimuli for four trial categories 00: 100, S0: 030, F0: 133 and FS: 131.

In all conditions, stimuli were horizontal rows of 3 digits presented in the center of the screen. Two of the digits were always identical, and the target digit was different from the other two. Digits 0, 1, 2 or 3 were used, but 0 was never a target and was not mapped to any of the response buttons. In control condition (00), a target digit value and its spatial position always matched each other and a target was flanked by two zeros (100, 020, 003). In Simon interference condition (S0), two zeros accompanied a target whose spatial position interfered with its value (010, 001, 200, 002, 300, 030). In the flanker condition (F0), value and spatial position of a target matched, but not-zeros flankers could evoke an erroneous response due to their mapping to response buttons (122, 133, 121, 323, 113, 223). Multi-source (flanker and Simon, FS) interference condition incorporated target value and position mismatch and response-linked flankers (212, 313, 221, 331, 211, 233, 112, 332, 131, 232, 311, 322).

Presentation software (Neurobehavioral Systems, Inc.) was used to program visual stimuli and to control the experiment. Stimuli were light gray (RGB: 226, 226, 226) and printed in Tahoma, and were presented on a dark background (RGB: 48, 48, 48). Viewing distance was kept to 55 cm to ensure that the height of visual stimulus had 1° and the length of the row had 3° visual angle (see (Bush et al., 2003) for guidelines). A scheme of experiment is shown in Figure 1. Each participant was presented with around 400 stimuli (approximately 100 stimuli per condition) separated in four runs with a 30 s break between runs. Within each run, two 10 s pauses were used to separate them into equal parts. Stimuli were presented in a mini-block design, with three to four stimuli of the same category per mini-block. The sequence of mini-blocks was pseudo-randomised with a restriction that the set of the same condition stimuli cannot repeat more than twice in a row over the course of the experiment. Each stimulus was presented for 900 ms and was followed by a blank screen lasting from 800 to 1300 ms (in 100 ms increments). A longer blank screen interval (2500-4400 ms) was placed after each mini-block, and a short break (10 s with white fixation cross on a screen) after 9-10 mini-blocks. In total, the experiment consisted of 4 runs (∼100 stimuli each), lasting approximately 22 minutes in total.

### 2.3 Behavioral data analysis

In-house MATLAB scripts were used for analysis of behavioral data. For each trial the reaction time (RT) and response type (1/2/3) were extracted from Presentation log files. Trials with no responses, multiple responses, incorrect responses, with RT shorter than 200 ms or longer than 1700 ms were classified as errors. Only correct trials were included in the analysis of RTs. Mean reaction time and accuracy (number of correct responses divided by the number of all trials per condition) were computed for each subject and task condition.

Statistical analyses of RTs and accuracy were performed using R (R Core Team, 2023) with RStudio (Posit team, 2022) self-written scripts. Used packages included: *tidyverse* (Wickham et al., 2019), *ggpubr* (Kassambara, 2023a), and *rstatix* (Kassambara, 2023b). ANOVA with repeated measures (within-subjects ANOVA) was conducted for normally distributed data, and replaced with the Friedman test when the data were not normally distributed. Distribution normality was controlled using the Shapiro-Wilk normality test and assumption of variance sphericity was confirmed with the Mauchly’s test of sphericity. In cases in which the assumption of sphericity was violated, the Greenhouse-Geisser correction was used. Post-hoc Student’s t-tests with Bonferroni correction (after ANOVA) or Wilcoxon signed-rank tests (after Friedman) were used to investigate differences between pairs of conditions. For analysis of interaction between conflict types (F0S0 vs. FS00) Student’s paired-samples t-test (for normally distributed data) or Wilcoxon signed-rank tests (for not-normally distributed data) was used.

### 2.4 EEG data acquisition

EEG data were collected with actiCHamp amplifier system (Brain Products, Munich, Germany) from 128 Ag-AgCl active electrodes (actiCAP, Brain Products, Munich, Germany) positioned on the elastic cap according to the International 10–5 system (Oostenveld and Praamstra, 2001, see Supplementary Fig. S1). TP9 and TP10 electrodes were placed on the left and right ear lobes, respectively, to enable alternative offline re-reference and were discarded from the analyses. Ground channel was set to FPz. EEG signal was on-line referenced to FCz electrode and digitized at 1000 Hz sampling rate with high cut-off filter set at 280 Hz. All impedances were kept below 10 kΩ (verified before and after completion of the task). At the end of the EEG session, we registered the exact positions of actiCap active electrodes on subjects’ heads in the 3D space by the use of CapTrak (Brain Products, Munich, Germany) hand-held scanner with two integrated cameras. EEG/ERP dataset (Dzianok et al., 2024) and its description (Dzianok et al., 2022) are freely available on-line.

### 2.5 Structural data acquisition

Anatomical MRI data were obtained on a 3T Siemens Prisma MRI scanner (Siemens Medical Solutions) using a 64-channel RF head coil, and with a T1-weighted 3D MP-Rage sequence (TR = 2.4s TI = 1000ms, TE = 2.74ms, 8° flip angle, matrix 320x320, FOV = 256x256 mm, isometric voxel size 0.8 mm, 240 slices), lasting 6:52 minutes (for more details see (Wojciechowski et al., 2024). Scans were obtained from 40 out of 41 subjects, one subject did not attend the scanning session and thus was excluded from the source analysis.

### 2.6 EEG data preprocessing

Brain Vision Analyzer software (Version 2.1.2.327, Brain Products, Munich, Germany) was used to preprocess EEG data. Artefacts (eye-movements, cardiac and muscle artefacts) were cleaned using Independent Component Analysis (ICA; Makeig et al., 1996). Excessive artefact-laden channels were replaced by using spherical splines interpolation (Perrin et al., 1989). EEG signals were band-pass filtered (0.5-40 Hz, order 8) and re-referenced to a common average reference. Remaining artefacts were marked and removed during final visual data inspection in Brain Vision Analyzer. Epochs with signal discontinuities (due to artefacts removal), bad responses (erroneous, too fast (earlier than 200 ms poststimulus), too slow (later than 1700 ms post-stimulus)), missing or multiple responses and with response from the previous trials falling into their baseline, were excluded from further analyses. On average, 24.30% (96.83±56.89) of trials per participant were excluded based on behavioral criteria and/or low signal quality.

For cluster based permutation ERP analyses, the preprocessed EEG data, event markers, and locations of the electrodes were imported from Brain Vision Analyzer to Matlab (R2018b) via EEGLAB (Delorme & Makeig, 2004) *bva-io* plug-in. Data was cut into 1100 ms epochs starting 200 ms before stimulus onset and baseline corrected.

For ERP topographic (microstate) analyses, artefact-free EEG data were segmented in Analyzer and averaged separately for each condition into four mean ERPs. No baseline correction was applied. Averaged ERPs from each participant were exported to text files and loaded to RAGU (Randomization Graphical User interface) software (Koenig et al., 2011).

Examples of obtained grand average ERPs from a few representative electrodes are presented in the Sup. Fig. S3. Note, that in the description of the waveforms (in the Supplementary Figure S3 and Result text) we stick to a nomenclature based on their latency and polarity.

### 2.7 Analysis of putative interactions of multi-source conflict effects

We tested if the effects of multiple conflicts presented concurrently sum up linearly, or whether there are non-linear effects which may indicate more complex interactions between underlying neural processes. For this purpose, we modelled the linear additive effect by combining trials recorded during flanker-only and Simon-only trials (F0 + S0; the number of trials included from each condition was equalised). This simulated effect was compared to the actual one observed in the combined conflicts condition (FS). Note that baseline effect was included in both Simon (baseline + Simon effect) and flanker (baseline + flanker effect) trials, thus to balance the equation, the no-conflict (00) condition had to be added to the multi-source one, so this modelled comparison can be expressed as FS00 *versus* F0S0.

### 2.8 Cluster based permutation ERP analysis

Stimulus-locked ERP analysis was performed using FieldTrip (Oostenveld et al., 2011). Trials were averaged separately for individual conditions and participants using *ft_timelockanalysis* function. The analysis started with testing for interaction between flanker and Simon conflicts, since its presence could confound main effects in 2x2 ANOVA design. To this means, we compared the additive model (F0S0) with actual activity recorded during simultaneous presence of multiple conflicts (FS00), revealing interaction between flanker and Simon effects. Thus, instead of computing main effects of each conflict type we computed simple effects of Simon (S0 vs. 00), flanker (F0 vs. 00) and MSIT (FS vs 00) interference. To reveal the differences between the two types of conflicts we also compared them directly with each other (F0 vs. S0).

Statistical comparison of multidimensional ERP data (126 channels x 1100 timepoints, from 200 ms before to 899 ms after stimulus onset) was performed using paired sample t-tests with cluster-based correction for multiple comparisons (Maris & Oostenveld, 2007) as implemented in *ft_timelockstatistics* function. Single-point statistics were thresholded at *p* = 0.01 (two tailed). Data-points with significant differences were clustered in space and time domains. To tighten the spatial extent of the obtained clusters, we specified that only samples with at least two neighboring channels expressing the same effect will be pooled together. Neighboring channels were defined based on the FieldTrip template file for our montage (*easycapM15_neighb*). Statistics of empirically obtained clusters (summed values of t-tests) were tested against the null distribution obtained in the 1000 permutations, with the significance threshold set at *p* = 0.05.

Since clusters defined by different comparisons have slightly different topographies and durations limiting direct comparisons between conditions, subsequently, we performed post-hoc paired comparisons of cluster magnitudes. For each trial type cluster magnitudes were calculated as follows: i) signal for all electrodes belonging to a cluster was averaged, ii) an integral was calculated of all the time points assigned to a cluster separately for positive and negative ERP waves (using *areap* and *arean* options from *getvalue* function as implemented in ERPLAB; (Lopez-Calderon & Luck, 2014), iii) magnitudes of simultaneously occurring clusters with the opposite polarity were considered as representing the same dipole-like effect, and were summed. To avoid assumption about distribution normality, post-hoc comparisons were run using permutation paired t-tests (with 2000 permutations). For each cluster we compared all pairs of conditions except the one which was used for cluster definition. To correct for multiple comparisons, the significance threshold was adjusted according to Bonferroni correction (i.e. corrected significance level for all comparisons p = 0.01, except comparison with simple additive model p = 0.0083).

### 2.9 Microstate analysis

Functional microstates are defined as topographic maps which remain stable for a short period of time, after which scalp field configuration rapidly changes from one microstate to another (Lehmann et al., 1987). Microstate analysis is based on the Global Field Power (GFP), i.e. standard deviation of the momentary potential values from all electrodes in a set (Lehmann & Skrandies, 1980), and the spatial distribution of these momentary potential values in scalp field maps (Koenig & Melie-García, 2009). ERP topographic analyses were conducted using the open-source RAGU (Randomization Graphical User interface) software (Koenig et al., 2011).

First, topographic consistency test (TCT; (Koenig & Melie-García, 2010) was applied to identify time windows with brain activity topographies consistent across participants for reliable microstate and source reconstruction analysis. TCT was applied with 5000 within-participant channel permutations, and p-values of topographic consistency were obtained for group-averaged waves for each condition at every time point. Additional testing for duration of significant TCT periods was applied and correction for multiple comparisons was provided as implemented in RAGU (Koenig & Melie-García, 2010). Only intervals with significant (p < 0.05) TCT were used in further analyses.

To cluster momentary ERP topographies into microstates, the k-mean clustering algorithm (Pascual-Marqui et al., 1995) was applied to the condition-specific, grand average ERPs from all 126 electrodes. Initial microstate maps were picked up randomly from local GFP maxima. Single-time-point topographies were then assigned to the initial maps which were continuously updated by averaging all momentary maps with the same label until explained variance became stable. The process was repeated 50 times with different initial maps and the final set of microstates was chosen as the one (from 50) with the highest percentage of explained variance. The optimal number of microstate cluster maps was first defined during cross-validation procedure (Koenig et al., 2014), during which the clustering (as described above) was performed with an increasing number of maps (3-12). ERPs from a randomly selected half of the subjects (training set) were used for clustering and the ERPs of the other half of the subjects (test set) were then fitted into obtained maps and the percent of explained variance was estimated. The analysis was performed for each of the tested map set sizes (3-12), and repeated 250 times with new random splits into training and test sets. The optimal number of microstates was selected as the one for which the average increment in the percent of explained variance was lower than 0.5% (see Supplementary Figure S1). The final set of microstate maps was computed using the whole available data and fitted to condition-specific (00, S0, F0, FS) grand average ERPs.

Values of all microstates’ parameters (onset, duration, area under the curve (AUC), offset, center of gravity and mean GFP were extracted to statistically analyze dynamic changes of ERP-microstates between conditions. The center of gravity provides information about the distribution of a GFP power within the microstate time-window. The AUC is a global measure of activation strength, considering both the duration and the GFP amplitude. It is measured as the sum of the GFP values from all the time samples (normalised by the sample duration [μV x ms]) where the selected microstate is present; the mean microstate’s GFP is computed as the average of these values.

Nine microstates from the optimal solution (see Supplementary Figure S1) were assigned to the grand mean ERPs of each condition (see Results 3.3), but further statistical microstate analysis was limited to those microstate classes that occurred during stimulus presentation (in the window 0-900 ms after stimulus onset). Microstates parameters were statistically compared between task conditions in 1 x 4 factorial design (factor condition: 00, S0, F0, FS). As noted above, the effects of coexistence of multiple conflicts were compared in a simple additive model in a 1 x 2 factorial design (factor condition: FS00 *versus* F0S0). Analysis was performed using a non-parametric randomization approach implemented in RAGU (Koenig et al., 2014) in which conditions were randomly assigned to individual ERPs within each subject dataset. The distribution of microstate parameters values under the null-hypothesis (distribution of effect happening by chance) was estimated from 5000 iterations of the procedure (the null-hypothesis was rejected at threshold of p < 0.05).

### 2.10 Source analysis

#### Forward model

The anatomically accurate forward volume conduction models were prepared individually for each participant using the FieldTrip-SimBio pipeline as described by (Vorwerk et al., 2018) with minor modifications, described below. First, head models were prepared using individual structural T1 images that were segmented using CAT12 toolbox (www.neuro.uni-jena.de/cat; Gaser et al., 2024). Six compartment types were identified: white matter (WM), gray matter (GM), cerebro-spinal fluid (CSF), skull (bone), eyes and the remaining soft tissue. Electrical conductivity assigned to segmented compartments was accordingly: 0.1429, 0.3333, 1.5385, 0.0063, 0.5000, 0.1000 S/m (see supplementary material in (Liu et al., 2018). Segmented images were downsampled to 2.0⨯2.0⨯2.0 mm voxel size in order to reduce computation time. Downsampled segmentations were used to construct hexahedral meshes with node shift set to 0.3 mm to obtain more smooth representations of tissue boundaries, as suggested by (Vorwerk et al., 2018). Potential sources within brain volume and its proximity (white matter, gray matter, CSF) were modeled using a Cartesian grid with 5 mm resolution. Digitized electrode locations were transferred to head geometry space using three fiducial points (i.e., nasion, left and right preauricular locations) using standard FieldTrip procedure followed by visual inspection (Oostenveld et al., 2011).

#### Linearly Constrained Minimum Variance Beamformer

Source localization was conducted on ERP data with pre-computed transfer matrix using linearly constrained minimum variance method (LCMV; Van Veen et al., 1997). LCMV uses covariance of signals recorded from the electrodes to generate spatial filters that maximize the activity from the individual source while simultaneously minimizing activity from all other sources. Common (i.e. for all conditions) covariance matrix was computed separately for each microstate. Since the latency and duration of microstates differed between conditions, to build a common covariance matrix the microstate time windows were cut to match the length of the shortest one. The reduction was applied equally at the onset and offset of a microstate. Covariance matrix was first calculated for individual trials and later averaged to minimize functional correlation between sources which could hinder their localization (Van Veen et al., 1997). Regularization of the covariance matrix was implemented and set to 5%. Source activity was expressed as an index of neuronal activity (NAI) as it is implemented in *ft_sourceanalysis*. Finally, estimated source power was interpolated into individual anatomical data.

#### Source level statistics

To enable group level analysis, anatomical images representing source activity were transformed to the standard MNI space using the new normalisation method as implemented in *ft_volumenormalise*. Normalized volumes representing source activity were then z-scored within each subject to ensure equal contribution of all subjects to group results, regardless of their average source power. Source level group statistics were performed using SPM12 software (Wellcome Trust Centre for Neuroimaging, UK). For each microstate one-sample t-test was conducted to reveal brain regions which activity was significantly increased. To effectively control for false positive results, obtained statistical maps were thresholded at *p* = 0.001 (voxel level; as recommended for similar analysis of fMRI data by Eklund et al., 2016). Clusters were considered significant if they reached *p* <= 0.05 FWE corrected. Anatomical description of significant clusters was obtained using Automatic Anatomical Labelling atlas (Rolls et al., 2020), and their overlap with large scale brain networks was determined by comparison with Cortical Functional Networks (Yeo et al., 2011) using MRIcron (MRIcron https://www.nitrc.org/projects/mricron) and applying obtained clusters of significant source activity as volume of interest. Maps were prepared in BrainNet toolbox (Xia et al., 2013), with nearest neighbor voxel interpolation and BrainMesh ICBM152 smoothed template. In a case of an atlas dataset, nearest neighbor interpolation produced a surface map with some discontinuities, which were removed (in Gimp v2.10.36) for better clarity of a figure. The original and edited maps can be seen in the Supplementary Figure S2. Next we computed the engagement of each functional brain network during subsequent microstates. To this means, for each microstate, we intersected group-level masks of source activity with masks of functional networks. Results of this intersection were normalized by the size of the network (to estimate engagement of each network at the given stage) as well as by the size of the source activation cluster (to expose importance of each network at the particular processing stage).

### 2.11 ERP microstates correlation analysis

Elongation of reaction times can reflect longer perceptual processing of the stimulus and/or differences in decision making and/or differences in response preparation and execution (Paluch et al., 2021). Toevaluate whether and how individual differences in performance were related to intensity and duration of specific processing steps, we conducted an analysis to test participant-wise correlations between the particular microstate features, duration, AUC and mean GFP, and behavioral measures. To evaluate whether the conflict processing steps were modulated by the preceding processes, correlation analyses between microstate features were performed for microstates that exhibited significant conflict-related changes in the group-level analysis (see Table 2) and the microstates directly preceding the microstates of interest. A similar approach was introduced by (Schiller et al., 2016), who applied it to explain individual differences in implicit bias. For each participant, individual mean ERPs for each condition were divided into microstate windows (using the same set of cluster maps as in the group-level analyses). Microstate parameters were extracted for each microstate and each subject. To account for individual differences in behavior unrelated to conflict-processing, RTs and accuracy in the non-conflict condition (00) were subtracted from those in the conflict trials (S0, F0, FS). The same standardization was applied to microstate features. We computed Spearman rank correlations and used the Benjamini-Hochberg procedure (Benjamini & Hochberg, 1995) to control for the False Discovery Rates and correct for multiple testing.

## 3. Results

### 3.1 Behavioral results

Analysis of RTs data revealed a gradual prolongation of responses from no-conflict (535.86±61.08 ms, mean±SD) through Simon (592.98±54.52 ms), flanker (676.85±68.82 ms) to multi-source (744.33±83.56 ms) conditions (Fig. 2A, F(1.72, 68.68) = 381.98, p < 0.001 with Greenhouse-Geisser correction). Post-hoc Student’s t-tests with Bonferroni correction indicated differences between all task conditions (all p < 0.001). Overall performance was high with only 4.30% (17.12±18.59) of trials qualified as errors. However, there were differences in accuracy across the task conditions (χ2(3, N = 41) = 74.90, p < 0.001). In line with RTs, the best accuracy was observed in 00 trials (proportion equal 0.98±0.04), and the worst for FS condition (0.88±0.15), with intermediate results for S0 (0.97±0.04) and F0 (0.96±0.07) conditions. Paired comparisons (Wilcoxon signed-rank test) showed that accuracy in Simon and flanker conditions was similar to each other (p = 0.80) but lower than in no-conflict condition (S0 vs. 00 p < 0.01, F0 vs. 00 p < 0.001). Performance in multi-source interference condition (FS) was lower than in all other conditions (all p < 0.001).

**Figure 2.**
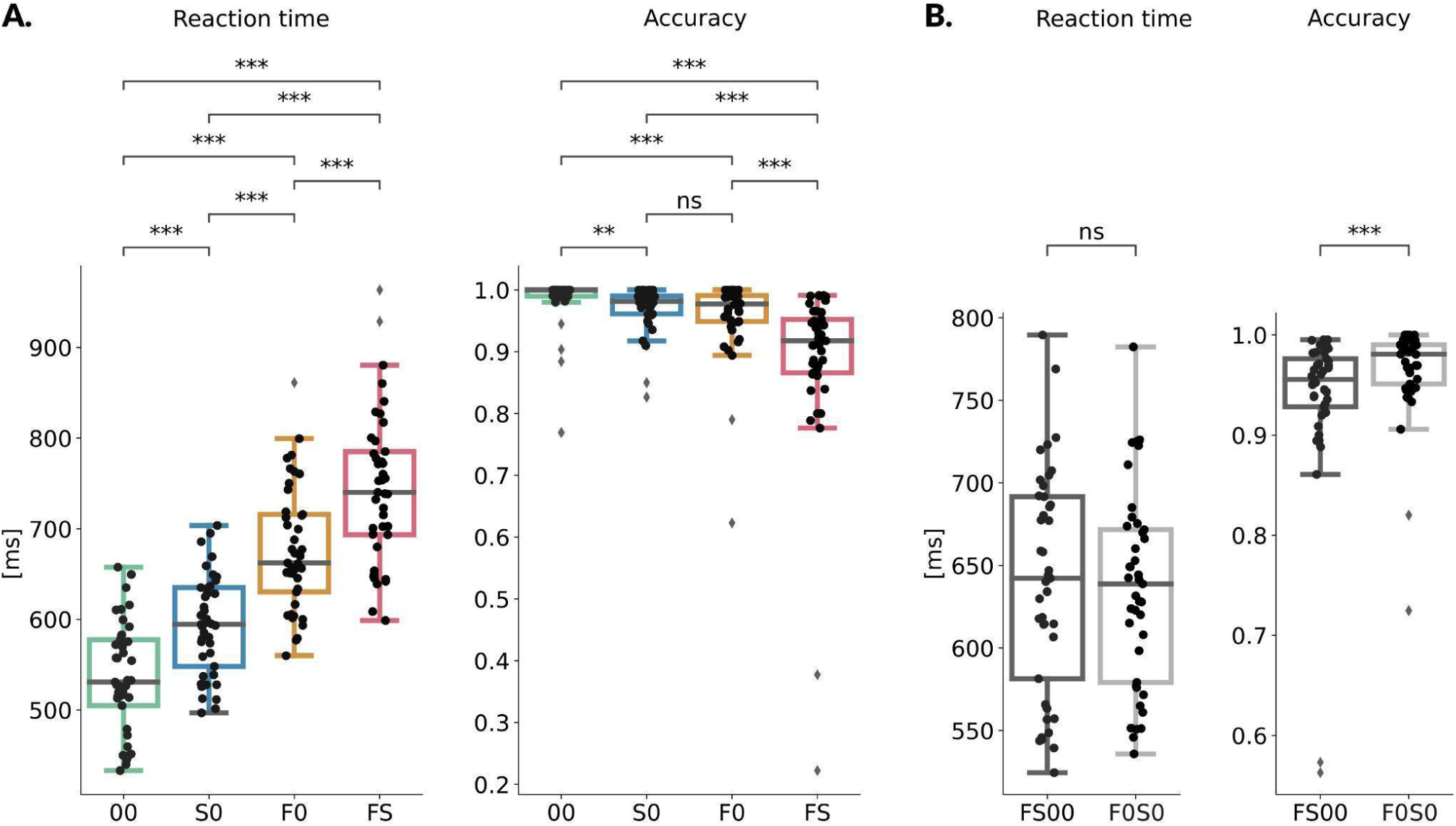
Behavioral results. A) Reaction time and accuracy for four MSIT conditions and B) for a comparison of a simple additive model (F0S0) with real multi-source data (FS00). Accuracy measured as a proportion of correct responses within all trials per condition. On each box, the central mark indicates the median, and the bottom and top edges of the box indicate the 25th and 75th percentiles, respectively. The whiskers extend to the most extreme data points, not considered outliers. Outliers are marked with diamonds. The significance of post hoc comparisons is marked with asterisks: *** p ≤ 0.001, ** p ≤ 0.01. Conditions are described in the X axis legend and marked by colors (note that the same color scheme is used in further figures presenting ERP results).

Finally, the effect of multi-source interference was compared to the additive model (FS00 vs F0S0, Fig.2B). This comparison showed that RTs between FS00 and F0S0 were similar (640.01±67.57 ms vs. 634.91±58.87 ms; t(40) = 1.76, p = 0.086). For accuracy, a significant difference was detected (Wilcoxon signed-rank test; z = 753, p < 0.001), with the concurrent presentation of the two conflicts characterized by slightly lower task performance (FS00 0.93±0.09), than expected from simple sum of the two effects presented in isolation (F0S0 0.96±0.05).

### 3.2 ERP waveforms results

Cluster-based permutation analysis of ERPs evoked by trials with Simon, flanker and multi-source interference, compared to no-conflict trials, identified three time windows during which (to various extent) brain activity was altered by the presence of conflict. The results are illustrated in Fig. 3 and Supplementary Figure S4, and detailed in Table 1 (cluster activity time windows, number of electrodes, and p-values) and Supplementary Table SI (p-values for post-hoc comparisons).

**Figure 3.**
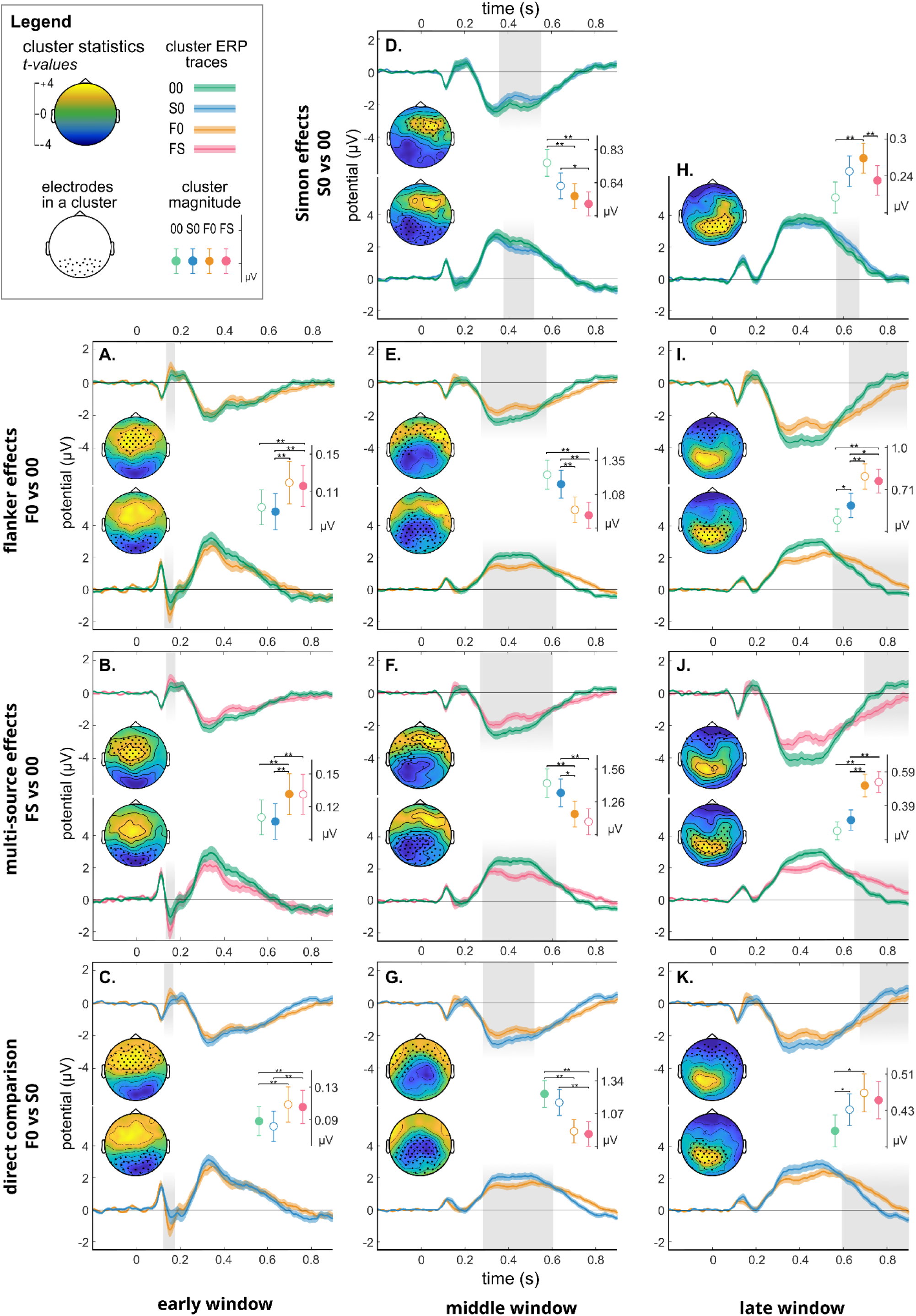
Results of cluster permutation analysis of ERP waveforms. Comparison of no-conflict trials with Simon (D, H), flanker (A, E, I) and multi-source (B, F, J) interference trials and a direct comparison of flanker and Simon trials (C, G, K). Topographic maps show the distribution of t-test values computed for the average (over the time of the significant difference) activity for each significant cluster (gray dots mark electrodes included in a cluster of significant difference). ERP traces represent averaged activity of all electrodes assigned to each cluster. Pale color areas along ERP trace represent ± SEM across subjects. Gray areas anchored on time axes mark time windows of significant differences defining duration of a particular cluster. Note that the activity maps are polarized in anterior-posterior axis and that most of the clusters form dipol-like pairs within similar time windows. Cluster magnitudes (area under the curve) from the time windows of significant differences are measured for each condition and plotted in inserted dot-charts (mean ±SEM); empty dots marked cluster-defining comparison, which was not included in post-hoc tests. Asterisks denote significance of post hoc comparisons equivalent to: *p < 0.05, **p < 0.01 after Bonferroni correction (see Methods).

**Table 1.** Results of cluster-based permutation ERP tests indicating the timing (onset, offset), number of involved electrodes (el. N) and p-values of detected significant differences between conflict and control (no-conflict) trials (see Suppl. Table SI for results of post-hoc comparisons).

|  |  | clusters with anterior topography |  |  |  | clusters with posterior topography |  |  |  |
| --- | --- | --- | --- | --- | --- | --- | --- | --- | --- |
|  |  | cluster-based permutation test result |  |  |  | cluster-based permutation test result |  |  |  |
|  |  | onset | offset | el. N | p-value | onset | offset | el. N | p-value |
| conflict versus no-conflict comparisons |  |  |  |  |  |  |  |  |  |
| early window<br>N150 / P150 | S0-00 | - | - | - | - | - | - | - | - |
|  | F0-00 | 132 ms | 172 ms | 39 | 0.029 | 126 ms | 169 ms | 30 | 0.016 |
|  | FS-00 | 135 ms | 179 ms | 44 | 0.028 | 132 ms | 173 ms | 26 | 0.032 |
| middle window<br>N330 / P330<br>and<br>N500 / P500 | S0-00 | 359 ms | 552 ms | 35 | <0.001 | 377 ms | 517 ms | 45 | <0.001 |
|  | F0-00 | 276 ms | 577 ms | 56 | <0.001 | 283 ms | 691 ms | 64 | <0.001 |
|  | FS-00 | 272 ms | 607 ms | 57 | <0.001 | 280 ms | 618 ms | 57 | <0.001 |
| late window<br>sustained wave over<br>550 ms | S0-00 | - | - | - | - | 565 ms | 671 ms | 28 | <0.001 |
|  | F0-00 | 625 ms | 893 ms | 33 | <0.001 | 550 ms | 899 ms | 47 | <0.001 |
|  | FS-00 | 696 ms | 899 ms | 23 | 0.003 | 650 ms | 899 ms | 45 | <0.001 |
|  |  | - | - | - | - | 737 ms | 862 ms | 8 | 0.028 |
| direct F0 versus S0 comparison |  |  |  |  |  |  |  |  |  |
| early window | F0-S0 | 126 ms | 169 ms | 60 | 0.005 | 122 ms | 174 ms | 37 | 0.007 |
| middle window |  | 282 ms | 518 ms | 50 | <0.001 | 282 ms | 604 ms | 61 | <0.001 |
| late window |  | 676 ms | 898 ms | 38 | <0.001 | 592 ms | 899 ms | 42 | <0.001 |

The earliest observed conflict-related effect (around 150 ms after stimulus onset) was specific to trials involving flanker interference (Fig. 3 A-C). It was manifested as an increased positivity over the frontal area (P150; F0 vs. 00: 132-172 ms, 39 electrodes, p = 0.029; FS vs. 00: 135-179 ms, 44 electrodes, p = 0.028; F0 vs. S0: 126-169 ms, 60, p = 0.005), and concurrent increased negativity over the occipital area (N150; F0 vs. 00: 126-169 ms, 30 electrodes, p = 0.016; FS vs. 00: 132-173 ms, 26 electrodes, p = 0.032; F0 vs. S0: 122-174 ms, 37, p = 0.007). Since clusters defined by different comparisons have slightly different topographies and durations, limiting direct comparisons between them, for each cluster we run post-hoc tests in which average magnitude (see Methods 2.8) was directly compared between conditions (see insets in Figure 3). Post-hoc comparisons confirmed that the N150/P150 modulation was specific to trials involving flanker conflict.

The subsequent effects were observed in the extended, complex waves of frontal negativity and posterior positivity spanning from around 250 to over 600 ms (Fig. 3D-G). These waves exhibited two local extrema (one around 330 ms, and the other around 500 ms), separated by a brief amplitude reduction around 420 ms. Compared to non-conflict trials, the amplitudes of both components (anterior-negative and posterior-positive) of this complex wave were reduced in the conflict trials. The effect was least pronounced for Simon only trials (S0 vs. 00; frontal negativity: 359-552 ms, 35 electrodes, p < 0.001; posterior positivity: 377-517 ms; 45 electrodes, p < 0.001; Fig. 3D), more pronounced for flanker only trials (F0 vs. 00; frontal negativity: 276-577 ms, 56 electrodes, p < 0.001; posterior positivity: 283-691 ms; 64 electrodes, p < 0.001; Fig. 3E) and the most pronounced in multi-source interference (FS vs. 00; frontal negativity: 272-607 ms, 56 electrodes, p < 0.001; posterior positivity: 280-618 ms; 57 electrodes, p < 0.001; Fig. 3F). A direct comparison of Simon and flanker interference revealed that, during this stage, the conflict-related reduction in amplitude was greater in flanker trials than in Simon trials (S0 vs. F0; frontal negativity: 282-518 ms, 50 electrodes, p < 0.001; posterior positivity: 282-604 ms; 61 electrodes, p < 0.001; Fig. 3G). Post-hoc comparisons, run separately for each cluster, showed that the reduction in amplitude due to Simon conflict was detectable only when topography and analysis window were defined by the Simon vs. non-conflict condition (Fig. 3D inset). However, this reduction was diluted when the analysis window was defined using other contrasts (insets in Fig. 3E-G). Overall, the amplitude reduction during this middle stage of stimulus processing was smaller in Simon trials compared to flanker and multi-source interference trials.

The final window of significant differences (Fig. 3H-K) fell on the falling slope of the above-mentioned long waveform and affected its both frontal-negative and posterior-positive components. In all comparisons, this late sustained potential (SP) had larger amplitude and/or longer duration in conflict than no-conflict trials. The least extensive effect was observed for S0 vs 00 comparison, where the differences were present in a short time window only on centro-parietal electrodes where the ERP wave had increased amplitude in S0 compared to 00 trials (S0 vs 00; frontal negativity: 565-671 ms, 28 electrodes, p < 0.001; Fig. 3H). Post-hocs paired comparisons revealed a strong effect also for F0 vs 00 but not for FS vs. 00 for which the area amplitude was significantly smaller than in F0 and not different from 00 (Fig. 3H, inset). In F0 vs 00 comparison (Fig. 3I), the increase in the amplitude of late sustained potential were much more extensive (F0 vs 00; frontal negativity: 625-893 ms, 33 electrodes, p < 0.001 and posterior positivity: 550-899 ms, 47 electrodes, p < 0.001). Cluster area amplitude was slightly increased in S0 trials and more increased in both types of trials including flanker-type conflict (Fig. 3I inset). In FS vs 00 comparison main clusters of significant differences started later but occupied similar topography as in case of F0 trials (FS vs 00; frontal negativity: 696-899 ms, 23 electrodes, p = 0.003 and posterior positivity: 650-899 ms, 45 electrodes, p < 0.001; Fig. 3J). Paired post-hoc results for FS vs 00 were similar as for F0 vs 00 (but with only trend level effect for S0 vs. 00 trials; Fig. 3J inset). There was also an additional, small (8 electrodes) cluster of FS vs 00 differences localized on the right temporo-parieto-occipital line (see Supplementary Figure S4 for average ERP waveforms and topography of the cluster). Direct comparison of Simon and flanker trials confirmed that conflict-related increase of ERP amplitude at this stage was more pronounced in flanker than in Simon trials (S0 vs F0; frontal negativity: 676-898 ms, 38 electrodes, p < 0.001 and posterior positivity: 592-899 ms, 42 electrodes, p < 0.001; Fig. 3K). To summarize, similar to the middle stage of the trial, all conflict types influenced ERP amplitudes during the late stage, however, duration and topography of this effect were conflict-specific. Moreover, post-hoc paired comparisons for Simon vs. no-conflict comparison suggested possible interaction of Simon and flanker conflicts. Specifically, the cluster area amplitude was greater in flanker-only trials compared to when flanker was combined with Simon interference (see Fig. 3H, inset). This effect was small and apparent only when the analysis window was defined by the Simon vs no-conflict comparison, which may explain why it has gone undetected in some of the previous studies (Frühholz et al., 2011b; Rey-Mermet et al., 2019b). We investigated it in detail in a separate analysis (see section *Interaction within the multi-source conflict*).

### 3.3 ERP microstates results

Visual inspection of ERP waveforms (Supplementary Figure S3) and results from the cluster-based analysis (Fig. 3) clearly indicated that the presence of different types of conflicts modulated not only the amplitude but also the temporal evolution of ERP responses. To better understand these effects, we applied microstate analysis, which is well-suited for detecting stable topographies of field potentials as well as task-related modifications in their duration and transitions. First, the consistency of topographies (TCT) was tested and confirmed across the entire analysis window (-200 to 1000 ms) for all four conditions (all p-values = 0.0002, corrected for multiple comparisons). The cross-validation procedure indicated that the set of nine microstates was optimal (Supplementary Figure S1); it explained 95.29% of variance in a full dataset. The topographic maps of these nine microstates and their assignments to the grand average GFP envelopes across the four experimental conditions are shown in Fig. 4.

**Figure 4.**
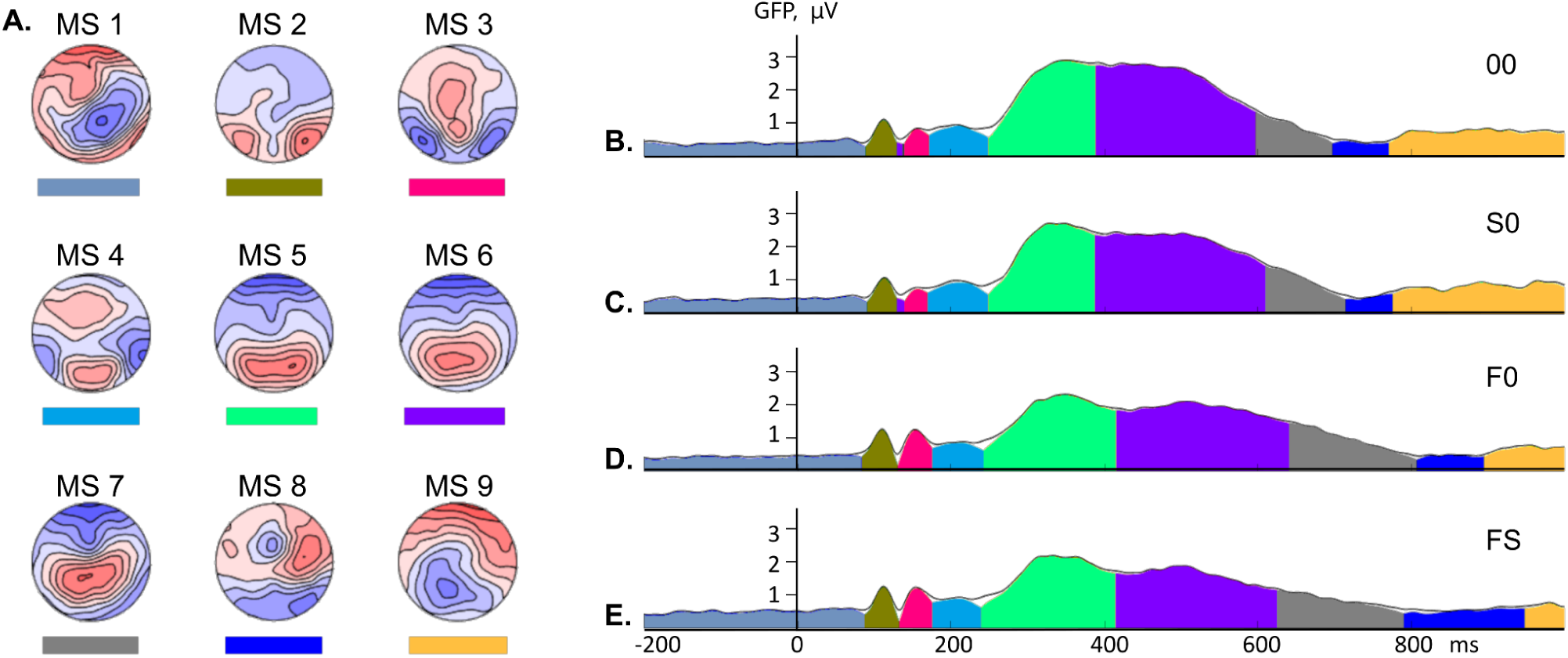
Microstate analysis results. A) The topographies of 9 microstates (MS 1-9). B-E) The grand average GFP envelopes for 00, S0, F0 and FS conditions. Individual microstate time-windows are marked by the color filing below the envelopes (color legend given in A, under each map) and the height of the colored areas under the black envelope line indicates the explained variance. Topographical consistency was high in the whole (-200-1000 ms) analytical window (TCT analysis, all p-values = 0.0002).

Despite the conflict-related differences revealed in the ERP cluster-based analysis, the number and sequence of observed microstates (further referred as MS1-9) were consistent across all conditions, including the no-conflict control condition. This indicates that no additional processes were evoked by the presence of single or combined interference, rather, the same processes were modulated differently depending on the type of conflict present (Fig. 4). MS1 was discarded from further analyses, as it represented baseline activity preceding stimulus onset. MS2 corresponded to the earliest exogenous ERP wave occurring 90-120 ms post-stimulus (occipital P110 and frontal N110). MS3 overlapped with the occipital negative and frontal positive waves, peaking around 150 ms (N150/P150 waves). The latency and topography of MS4 were similar to the temporo-parietal N200 (accompanied by two midline patches of positivity). Interestingly, the later, long complex wave of fronto-central negativity and parieto-occipital positivity was, based on topographical differences, divided into a few consecutive microstates (MS5, MS6, MS7). MS5 was characterized by a broad posterior positivity centered around 330 ms and counterbalanced by a frontal negativity (P/N330). In MS6, parietal positivity shifted to the midline around 500 ms after stimulus onset (P/N500). During the late sustained phase of this wave, this positivity moved towards the vertex, forming the MS7 map (centered around 650 ms after stimulus onset). The late sustained potential concluded with a brief MS8, characterized by hemispheric asymmetry. Finally, MS9 coincided with stimulus offset and predominantly occurred after task execution, and was therefore excluded from further analyses.

To capture conflict-related effects we measured different microstate features (e.g. onset, offset, duration, AUC; for details see Methods 2.9) and compared them across conditions (see Table 2). Note that there is an interplay between these features (e.g. AUC depends on duration and mean GFP) thus, instead of focusing on single features we will rather discuss their combinations. Overall, flanker interference influenced microstate dynamics more profoundly and earlier than Simon interference. The flanker-specific effect observed in ERP cluster-based analysis at the early stage of stimulus processing was closely reflected by MS3, which began earlier, lasted longer, and was more intense (measured with AUC and mean GFP) in trials including flaker conflict compared to the Simon and no-conflict conditions. However, the middle and late stage effects were proved to be more complex in the microstate analysis, which differentiated early P3-like map (MS5), late P3-like map (MS6) and divided the late sustained potential into two consecutive maps (MS7 and MS8). MS5 had lower intensity (measured by mean GFP) but lasted longer in trials with flanker conflict (F0, FS) compared to the no-conflict and Simon trials. In consequence, the following microstates were postponed (had later onsets) when flanker conflict was present compared to the no-conflict and Simon trials. The prolongation of MS5 compensated for its lower intensity as its overall power (measured with AUC) was similar in all compared conditions. In contrast, MS6 had similar duration across conditions accompanied with a smaller intensity (measured with GFP) which resulted in overall reduced power of this microstate in the presence of flanker interference. In the flanker-containing trials, MS7 started slightly later, and lasted longer than in no-conflict trials (significantly later center of gravity and offset). This prolongation led to a significant increase of microstate power (AUC) in flanker-only trials compared to no-conflict conditions but not when flanker was combined with Simon (FS vs. 00). The final, eighth microstate was significantly later (as indicated by measures of onset, offset and center of gravity) in all trials including flanker conflict and in FS its duration was longer, leading to stronger AUC than no-conflict condition. There were no Simon-specific effects (in comparison to no-conflict trials) for early (MS2-MS4) microstates. Slight modulation was observed for later processes, including lower intensity (mean GFP) of MS5 and MS6 and a tendency for stronger MS7 (trend level increase of mean GFP and later offset; both resulting in slightly, but significantly later center of gravity). Consequently, there was also a tendency for a final microstate (MS8) to be slightly later (trend level effects for onset and center of gravity) in Simon compared to non-conflict trials.

**Table 2.** Comparison of microstate features between different types of trials. Note that parameters of MS6 (in 00 and, S0 and FS conditions) and MS7 (in 00 condition) were measured without taking into account their short intrusions between MS2 and MS3. Significant p-values (<0.05) are written with bold font, trend-level (<0.1) with italic font and non-significant values in gray.

| MS class | microstate measures |  |  |  |  |  | statistical comparisons p-values |  |  |  |  |
| --- | --- | --- | --- | --- | --- | --- | --- | --- | --- | --- | --- |
|  | Microstate feature | 00 | S0 | F0 | FS | [units] | anova | S0-00 | F0-00 | FS-00 | S0-F0 |
| MS2 | Onset | 92.0 | 94.0 | 86.0 | 91.0 | ms | 0.384 | 0.685 | 0.311 | 0.946 | 0.130 |
|  | Duration | 39.0 | 37.0 | 46.0 | 41.0 | ms | 0.267 | 0.693 | 0.187 | 0.540 | *0.095* |
| | AUC | 29.9 | 29.6 | 38.3 | 35.4 | ms*μV | 0.136 | 0.924 | 0.096 | 0.189 | *0.063* |
|  | Offset | 130.0 | 130.0 | 131.0 | 131.0 | ms | 0.420 | 1.200 | 0.777 | 0.193 | 0.802 |
|  | Center of gravity | 112.0 | 112.8 | 108.7 | 111.6 | ms | *0.088* | 0.572 | 0.105 | 0.947 | **0.028** |
| | Mean GFP | 0.809 | 0.849 | 0.879 | 0.916 | μV | 0.423 | 0.387 | 0.224 | 0.138 | 0.585 |
| MS3 | Onset | 141.0 | 142.0 | 132.0 | 135.0 | ms | **<0.001** | 1.200 | **0.003** | **0.017** | **<0.001** |
|  | Duration | 32.0 | 29.0 | 45.0 | 42.0 | ms | **<0.001** | 0.702 | **<0.001** | **<0.001** | **<0.001** |
| | AUC | 22.0 | 19.0 | 40.9 | 39.6 | ms*μV | **<0.001** | 0.480 | **<0.001** | **<0.001** | **<0.001** |
|  | Offset | 172.0 | 170.0 | 176.0 | 176.0 | ms | **<0.001** | 0.624 | **0.033** | *0.055* | **0.004** |
|  | Center of gravity | 157.3 | 156.7 | 155.8 | 157.0 | ms | 0.782 | 0.518 | 0.332 | 0.810 | 0.703 |
| | Mean GFP | 0.752 | 0.745 | 0.983 | 1.015 | μV | **<0.001** | 0.827 | **<0.001** | **<0.001** | **<0.001** |
| MS4 | Onset | 173.0 | 171.0 | 177.0 | 177.0 | ms | **<0.001** | 0.624 | **0.033** | *0.055* | **0.004** |
|  | Duration | 77.0 | 79.0 | 67.0 | 64.0 | ms | **<0.001** | 0.668 | **0.021** | **0.009** | **0.004** |
| | AUC | 59.6 | 64.4 | 51.6 | 51.4 | ms*μV | **<0.001** | 0.138 | **0.023** | **0.044** | **<0.001** |
|  | Offset | 249.0 | 249.0 | 243.0 | 240.0 | ms | **0.031** | 1.000 | 0.158 | *0.065* | *0.099* |
|  | Center of gravity | 210.3 | 210.0 | 209.7 | 207.8 | ms | 0.494 | 0.831 | 0.756 | 0.277 | 0.861 |
| | Mean GFP | 0.858 | 0.892 | 0.841 | 0.883 | μV | 0.257 | 0.213 | 0.422 | 0.377 | *0.094* |
| MS5 | Onset | 250.0 | 250.0 | 244.0 | 241.0 | ms | **0.031** | 1.200 | 0.158 | *0.065* | *0.099* |
|  | Duration | 140.0 | 138.0 | 173.0 | 174.0 | ms | **<0.001** | 0.883 | **0.002** | **0.003** | **<0.001** |
| | AUC | 309.7 | 289.5 | 311.0 | 296.8 | ms*μV | 0.712 | 0.435 | 0.961 | 0.592 | 0.249 |
|  | Offset | 389.0 | 387.0 | 416.0 | 414.0 | ms | **<0.001** | 0.858 | **0.008** | **0.013** | **<0.001** |
|  | Center of gravity | 331.6 | 329.4 | 339.7 | 336.9 | ms | *0.073* | 0.628 | 0.126 | 0.308 | **0.014** |
| | Mean GFP | 2.266 | 2.147 | 1.840 | 1.753 | μV | **<0.001** | **0.044** | **<0.001** | **<0.001** | **<0.001** |
| MS6 | Onset | 390.0 | 388.0 | 417.0 | 415.0 | ms | **<0.001** | 0.858 | **0.008** | **0.013** | **<0.001** |
|  | Duration | 204.0 | 218.0 | 223.0 | 208.0 | ms | 0.721 | 0.180 | 0.709 | 0.945 | 0.793 |
| | AUC | 490.4 | 482.8 | 411.5 | 328.4 | ms*μV | **<0.001** | 0.806 | 0.115 | 0.001 | *0.082* |
|  | Offset | 593.0 | 605.0 | 639.0 | 622.0 | ms | **0.046** | 0.236 | **0.007** | 0.268 | **0.016** |
|  | Center of gravity | 480.5 | 487.0 | 524.8 | 511.6 | ms | **<0.001** | 0.246 | **<0.001** | **0.005** | **<0.001** |
| | Mean GFP | 2.370 | 2.141 | 1.864 | 1.581 | μV | **<0.001** | **0.003** | **<0.001** | **<0.001** | **0.002** |
| MS7 | Onset | 594.0 | 606.0 | 640.0 | 623.0 | ms | **0.046** | 0.236 | **0.007** | 0.268 | 0.382 |
|  | Duration | 101.0 | 104.0 | 163.0 | 160.0 | ms | **0.004** | 0.959 | **0.035** | 0.156 | **<0.001** |
| | AUC | 93.4 | 107.3 | 154.3 | 133.2 | ms*μV | *0.086* | 0.322 | **0.031** | 0.259 | **0.020** |
|  | Offset | 694.0 | 709.0 | 802.0 | 782.0 | ms | **<0.001** | **0.084** | **<0.001** | **<0.001** | **<0.001** |
|  | Center of gravity | 628.0 | 648.7 | 705.2 | 692.3 | ms | **<0.001** | **0.014** | **<0.001** | **0.002** | **<0.001** |
| | Mean GFP | 0.933 | 1.066 | 0.969 | 0.868 | μV | 0.320 | *0.062* | 0.811 | 0.642 | 0.294 |
| MS8 | Onset | 695.0 | 710.0 | 803.0 | 783.0 | ms | **<0.001** | *0.084* | **<0.001** | **<0.001** | **<0.001** |
|  | Duration | 53.0 | 55.0 | 81.0 | 119.0 | ms | **0.007** | 0.905 | 0.172 | **0.029** | 0.188 |
| | AUC | 21.2 | 26.0 | 34.9 | 58.1 | ms*μV | **0.007** | 0.420 | 0.116 | **0.018** | 0.373 |
|  | Offset | 747.0 | 764.0 | 883.0 | 901.0 | ms | **<0.001** | 0.266 | **<0.001** | **<0.001** | **<0.001** |
|  | Center of gravity | 719.6 | 737.3 | 843.0 | 842.0 | ms | **<0.001** | *0.064* | **<0.001** | **<0.001** | **<0.001** |
| | Mean GFP | 0.447 | 0.508 | 0.462 | 0.531 | μV | 0.442 | 0.162 | 0.772 | 0.197 | 0.459 |

### 3.4 Interaction within the multi-source conflict

The test for interaction between flanker and Simon conflicts was defined as a contrast of the activation during multi-source interference trials *versus* the additive model of two single-conflict trials (FS00 vs. F0S0, see Methods 2.7). Brain activity during the concurrent presentation of Simon and flanker interference was lower than expected from the additive model. In the ERP analysis (Fig. 5, Table 3A), the effect occurred during the slow potential wave, with multi-source interference trials (FS00) showing reduced (in comparison to F0S0 model) frontal negativity (541-724 ms, 40 electrodes, p < 0.001) and reduced centro-parietal positivity (524-733 ms, 36 electrodes, p < 0.001). To explore this effect further, we compared the magnitude of the identified cluster in all four trial types (inset in Fig. 5A). Both conflict types, when presented separately, increased the slow potential amplitude, with a slightly (but non-significantly) greater increase in response to flanker than Simon conflict. However, when the two conflicts were presented concurrently (FS), the magnitude of slow potential was not different from the no-conflict condition and was significantly lower than in flanker-only trials (Fig. 5A inset).

**Figure 5.**
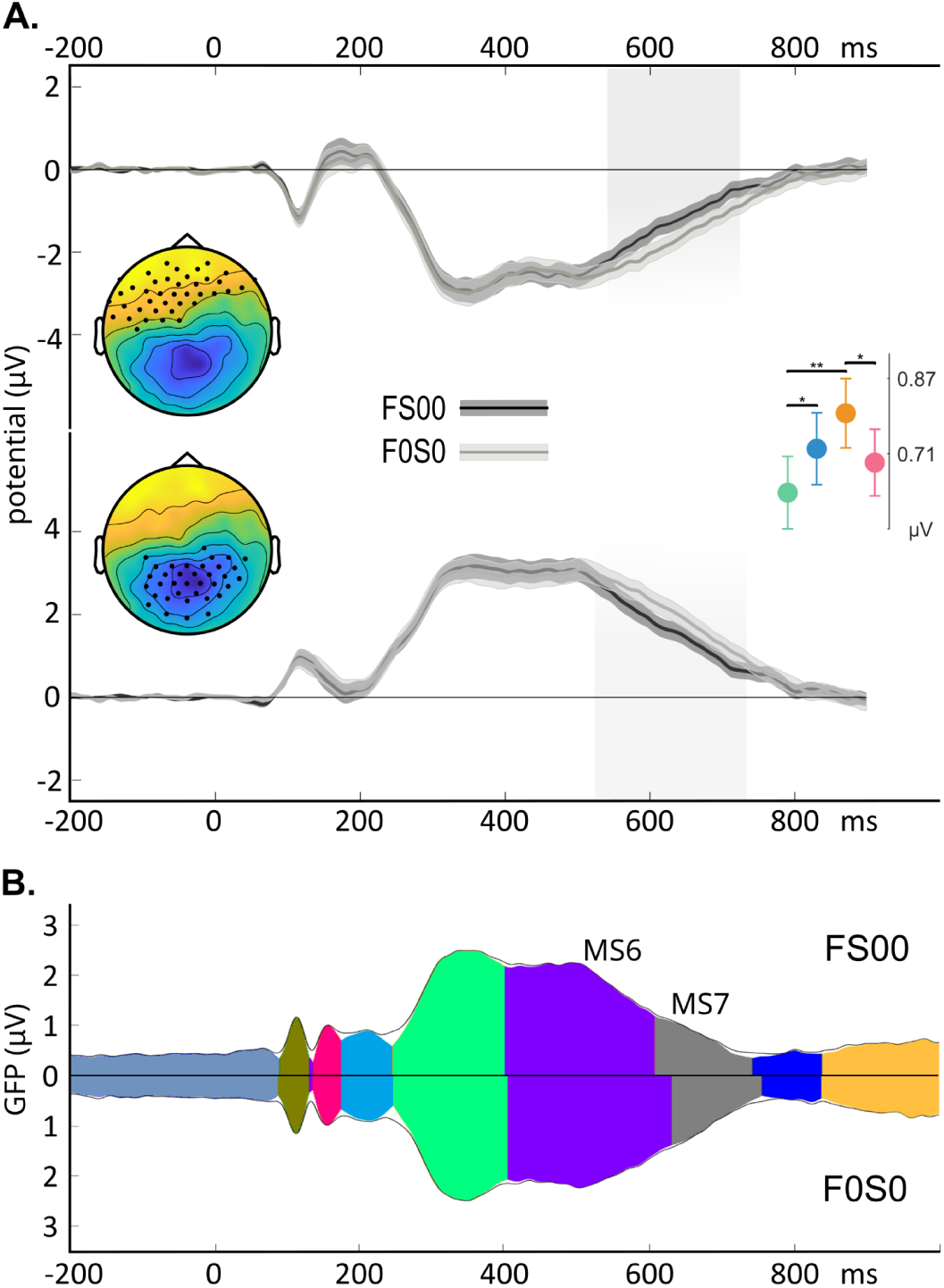
Interaction effect in multi-source conflict trials. A) Cluster based ERP analysis detected two corresponding clusters of difference between multi-source conflict (FS00, black) and additive model (F0S0, gray) data (post-hoc chart legend: green - 00; blue - S0; orange - F0, pink - FS). B) Microstates analysis – ERP global field power (GFP) envelopes for multi-source conflict data (FS00) and for additive model estimation (F0S0) counterposed along the X axis. Individual microstate time-windows are marked by the color filing envelopes and the height of the colored areas under the black envelope line indicates the explained variance (for microstate topographies and color legends see Figure 4A). Significant differences were detected for MS6 (violet) and MS7 (gray), for the numerical data see Table 3.

**Table 3.** ERP clusters and microstate features – interaction

| ERP cluster-based permutation test |  |  |  |  |  |
| --- | --- | --- | --- | --- | --- |
| late window | cluster topography: |  | anterior |  | posterior |
|  | Onset | ms | 541.0 |  | 524.0 |
|  | Offset | ms | 724.0 |  | 733.0 |
|  | electrodes number |  | 40 |  | 36 |
|  | p-value |  | <0.001 |  | <0.001 |
| post-hoc comparisons of cluster magnitudes |  |  |  |  |  |
| p-values (Bonferroni corrected significance threshold = 0.0083) |  |  |  |  |  |
| S0-00 | F0-00 | FS-00 | F0-S0 | FS-S0 | FS-F0 |
| 0.005 | < 0.001 | 0.179 | 0.113 | 0.565 | 0.005 |
| microstate analysis |  |  |  |  |  |
| MS class | microstate measures |  |  |  | p-value |
|  | feature | unit | FS00 | F0S0 |  |
| MS6 | Onset | ms | 401.0 | 405.0 | 0.541 |
|  | Duration | ms | 207.0 | 226.0 | 0.184 |
|  | AUC | ms*μV | 406.4 | 447.7 | 0.069 |
|  | Offset | ms | 607.0 | 630.0 | 0.046 |
|  | Center of gravity | ms | 493.6 | 510.0 | 0.005 |
|  | Mean GFP | μV | 1.932 | 1.960 | 0.540 |
| MS7 | Onset | ms | 608.0 | 631.0 | 0.115 |
|  | Duration | ms | 135.0 | 125.0 | 0.492 |
|  | AUC | ms*μV | 103.3 | 112.4 | 0.592 |
|  | Offset | ms | 742.0 | 755.0 | 0.339 |
|  | Center of gravity | ms | 661.3 | 680.6 | 0.048 |
|  | Mean GFP | μV | 0.800 | 0.924 | 0.046 |

The same microstate maps, occuring in the same sequence, were detected for the additive model as for the multi-source interference (Fig. 6B), but the features of MS6 and MS7 captured the interaction effect (Fig. 5B and Table 3B). MS6 showed an earlier offset and center of gravity and a trend for lower AUC in multi-source interference trials than predicted by the additive model. Similarly, MS7 was earlier (center of gravity parameter) and weaker (mean GFP) in multi-source trials than expected by the additive model. In summary, the late processing stages (MS6 and MS7) were less intense when conflicts were presented concurrently.

**Figure 6.**
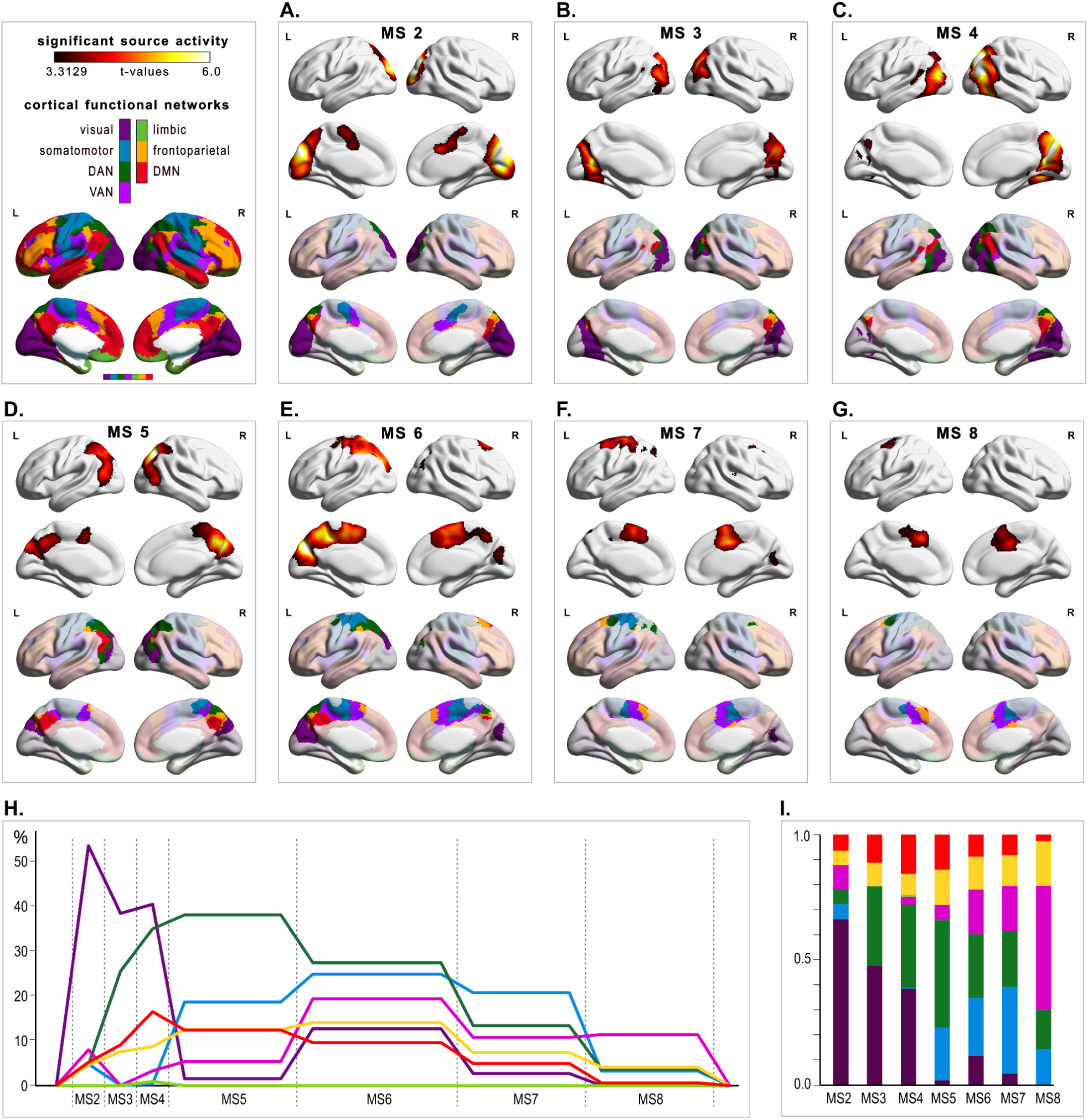
Brain sources of subsequent microstates and their correspondence to functional networks atlas. A-G) Brain activity maps: the upper part of each panel presents surface maps of significant source activity clusters; the lower part displays network maps masked according to the corresponding source activation. H-I) Schematic summaries of network dynamics and contributions: H) network activations normalized by the size of each network (to estimate engagement of each network during subsequent microstates). The X axis is arbitrarily adjusted according to the duration of microstates for FS condition; I) network activations normalized by the size of the source cluster (proportion of active networks contributing to the brain sources of each microstate is depicted). The brain network color legend applies to all panels.

### 3.5 Source reconstruction results

Microstate analysis allowed us to divide ERPs into time windows with stable topographies, ensuring the semi-stationarity of the associated brain processes. This in turn allowed us to apply the beamformer technique to localize the brain sources underlying each microstate (from MS2 to MS8). The identified sources were then anatomically described using the Automatic Anatomical Labeling (AAL) atlas (Rolls et al., 2020) and compared with the Cortical Functional Networks Atlas (Yeo et al., 2011). Surface projections of the sources and their overlay with Yeo’s Atlas are presented in Figure 6. Supplementary Table SII provides detailed information on the anatomical composition, values of statistical tests of the localized sources, including their size and MNI coordinates of peak activity. During our task we observed activation of various parts of all (except limbic) functional large scale brain networks, which were previously identified in resting state fMRI studies (Yeo et al., 2011): visual, somatomotor, dorsal attention network (DAN), ventral attention network (VAN), frontoparietal (FPN), default mode network (DMN). Lack of limbic activity is likely due to the type of task material (digits) which did not engage emotional processing. Importantly, contributions of different networks dynamically changed in subsequent stages of the task (see Fig. 6H-I).

The earliest analyzed microstate (MS2, ERP waves P/N 110 ms) was associated with increased activity in the occipital and medial frontal cortex (Fig. 6A, top panel). Comparison with functional networks revealed that the occipital cluster of activity primarily involved the visual network (Fig. 6A bottom panel). A second cluster of activation during MS2 was localized in the medial frontal cortex, encompassing the middle cingulate and paracingulate gyri, and extended into the supplementary motor areas (SMA) and to the left paracentral lobule (see Supplementary Table SII). These activations overlapped with the frontal node of the ventral attention network (VAN) and somatomotor network (reflecting movement related activation; Fig. 4A, bottom panel).

The sources of MS3 indicated the progression of information upstream the visual system towards the parietal lobe. The localization of MS3 sources primarily overlapped with higher visual areas and posterior nodes of the dorsal attention network (DAN), specifically the right and left intraparietal sulcus (Fig. 6B). Further upstream information flow was evident in the sources of MS4, with increased engagement of posterior nodes of DAN and the default mode network (DMN), visible as engagement of the area near the right angular gyrus. At this stage, overall source activity was stronger in the right hemisphere than in the left (Fig. 6C).

Source analysis for MS5 detected a cluster of activity over the dorsal occipital and parietal cortices (Fig. 6D, top panel). At this stage (corresponding to P/N330) activation, for the second time, reached the motor system (including left paracentral lobule and both postcentral gyri, see Supplementary Table SII), which remained active until the end of trial execution. Similarly to the medial frontal cluster observed during MS2, MS5 brain sources included anterior areas, such as the left middle cingulate and paracingulate gyri and the left supplementary motor area. Comparison with functional brain networks showed that parietal cortex activation aligned with the posterior nodes of DAN (i.e. intraparietal sulcus), activation in the vicinity of angular gyrus corresponded to parts of DMN and frontoparietal network (FPN) and activation over the medial frontal cortex reflected the engagement of VAN (Fig. 6D, bottom panel).

The sources of MS6 were similar to those of MS5 within the occipital and parietal cortices but extended over a significantly larger portion of the medial frontal cortex (reflecting involvement of the somatomotor system as well as VAN and FPN; Fig. 6E). Activity during this time was more prominent in the left hemisphere (likely related to the preparation and/or execution of right-hand movements). In addition to the regions of the occipital and parietal cortices already activated during the preceding microstate, MS6 was characterized by activation of the left angular gyrus (corresponding to the part of temporoparietal junction belonging to FPN), areas near the primary motor cortex, and supplementary motor area. Activation within the medial frontal cortex extended over the middle cingulate and paracingulate gyri (see Supplementary Table SII).

The final two microstates, MS7 and MS8, showed similar source activations concentrated in the medial frontal cortex and small areas of the dorsal frontal cortex. During MS7, most of the activity was localized in a frontal cluster encompassing motor areas (with peak activation within the primary motor cortex) and the medial frontal cortex encompassing the supplementary motor areas extending to the middle cingulate and paracingulate gyri (see Fig. 6F and Supplementary Table SII). Comparison with functional brain networks revealed engagement of somatomotor network, VAN and FPN (in the left hemisphere). Dorsal frontal cortex activations were localized in the superior frontal gyri and overlapped with the anterior nodes of DAN (i.e. frontal eye fields). A small occipital cluster was restricted to the right hemisphere and belonged to the visual network.

During MS8, one cluster of increased activity was detected, with a peak localized within the left supplementary motor area. Activations at this stage involved regions similar to those during MS7, though with less prominent activation of the motor cortex. Activation in the medial frontal cortex was primarily associated with VAN and FPN (Fig. 6G).

In summary, during the early stage of the trial, up to approximately 250 ms after stimulus onset (corresponding to microstates MS2-MS4), the majority of recorded brain activity originated from the visual network (Fig. 6H). This was followed by activation of DAN, which began (during MS3) in its posterior nodes and propagated upstream, peaking during MS5 (corresponding to P3a), approximately 250-400 ms after stimulus onset (Fig. 6H). The ventral attention network, the other of the two large attention systems, showed dynamics opposite to those of DAN. Its medial frontal node was transiently activated at the start of the trial, remained inactive during the visual processing and target selection stages, and reactivated around 388-417 ms after stimulus onset (MS6), coinciding with decline in activation of DAN’s posterior nodes (Fig. 6H). Interestingly, a similar pattern was observed in a somatomotor network, represented by activation of paracentral lobule, precentral gyrus and supplementary motor area. The last two networks which were engaged during our task were FPN and DMN. Their activation was modest (FPN was maximally activated at about 12% of its total volume, DMN at about 16%) but consistent throughout the trial. However, DMN was more involved during stimulus-related stages (similarly to DAN), while FPN was more engaged during response-related stages (similarly to VAN; see Fig. 6H,I).

### 3.6 Correlations of microstate features and behavioral differences

In our final analysis, we looked for correlation between task related changes of microstate features and performance. Correlation plots, coefficient values and significance levels are presented in Fig. 7. Behavioral cost of Simon conflict (increase in RT compared to no-conflict control condition) was, between-subjects, correlated with the change in the intensity of MS6 (difference in mean GFP compared to the non-conflict control). The more intensified was MS6 the smaller were behavioral costs of Simon (Fig. 7A, rho = -0.464, p = 0.002). Analogical relation was observed for flanker but with the intensity of MS7 (Fig. 7B, RTs, rho = -0.422, p = 0.006). Behavioral costs of flanker conflict were paralleled also by the prolongation of MS5 which correlated, between-subjects, with the accuracy drop (rho = -0.383, p = 0.013, Fig. 7C). The correlation analysis also revealed that, in the flanker task, the prolongation of the MS5 was preceded by a shortening of MS4 (Fig. 7D, rho = -0.437, p = 0.004). No other significant correlations between features of subsequent microstates were observed.

**Figure 7.**
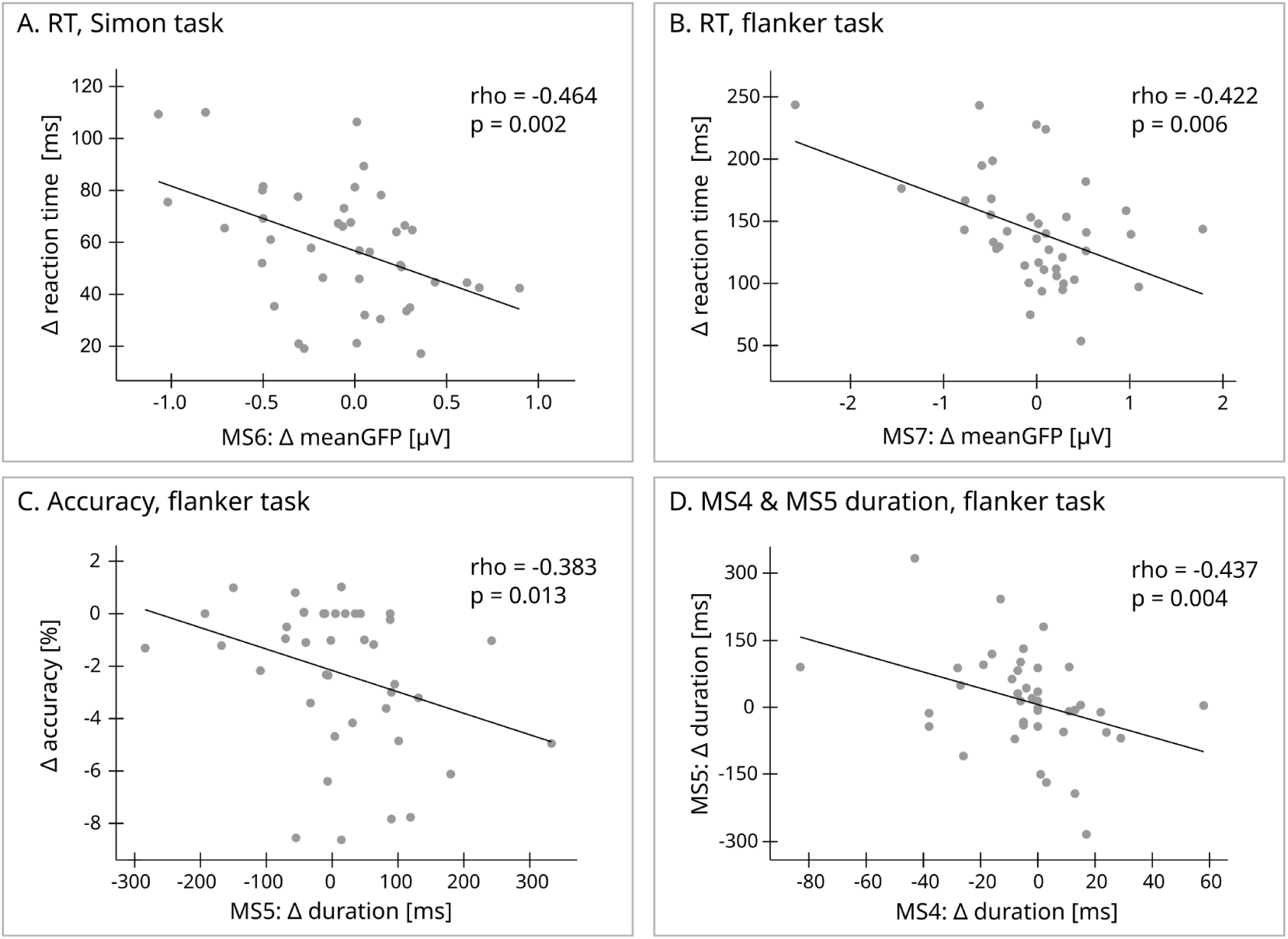
Correlation analysis of subsequent microstates features and microstate features and behavioral differences. A-B. Longer RTs were correlated with weaker (lower GFP) processing during MS6 (in Simon, S0 trials) and MS7 (in flanker, F0 trials); C. Worse accuracy in flanker trials was preceded by a longer MS5; D. Longer MS5 followed shorter MS4 in flanker trials. The values used in the analysis and plotted in the figure are the differences (as denoted by a Δ sign) from control, no-conflict condition (00 trials). Each dot represents one participant. Solid lines represent best fitted linear trends.

## 4. Discussion

In this study, we explored the spatio-temporal dynamics of uni- and multi-source conflict processing using the extended version of the multi-source interference task with four conditions, allowing us to compare various conflicts within the same stimuli set and a single experimental session. Exactly the same set of 3-digit stimuli were used in a parallel fMRI experiment conducted with the same group of participants (Wojciechowski et al., 2024). Behavioral data confirmed that our task was efficient in eliciting conflict, as evidenced by a gradual prolongation of RTs and increase in error rates from no-conflict, through Simon, flanker to multi-source conflict conditions (see Fig.2). Instead of focusing on pre-defined waveforms, we employed cluster permutation tests on ERP waveforms recorded from all sites during the entire trial duration, to identify conflict-sensitive effects. Subsequently, we used microstate analysis to delineate time windows corresponding to distinct processing stages. We applied beamformer source localization, to map the subsequent processing stages onto large-scale brain network activations identified in the resting state connectivity fMRI studies (Yeo et al., 2011).

### 4.1 Common processing stages are activated in all task conditions but with different dynamics

Analysis of ERP waveforms revealed three time windows which were affected by the presence of different types of conflicts. The early time window centered around 150 ms after stimulus onset, the middle time window ranging from 200 to 600 ms and the late time window from 600 to 800 ms after stimulus onset. Only the early effect was specific to flanker-type conflict while the two remaining effects were modulated by all conflict types but with conflict-specific differences in intensity and durations of the effects. Mirroring increasing costs of conflict observed in our behavioral data (see Fig. 2) effects of conflict presence on ERP were smallest for Simon, more pronounced for flanker and the most prominent for multi-source trials (see Fig. 3 and Table 1). Microstate analysis confirmed that ERP responses to control (no-conflict), Simon, flanker and multi-source interference stimuli all consisted of seven common processing stages, represented by consecutive microstates (MS2-MS8), occurring in the exact same order regardless of trial type. No additional processes were evoked by the presence of single or combined interference. The observed differences in ERP-microstate features were quantitative rather than qualitative. Trial types differed in the latencies, durations, and intensities of successive microstates (see Fig. 4 and Table 2).

Our findings align with previous ERP studies (Ruggeri et al., 2019; Schiller et al., 2016). Despite using different tasks to induce conflict – implicit association task (IAT) and Stroop color-word task – both studies reported no differences in the number and sequence of processing stages between congruent and incongruent condition. Instead, the presence of conflict resulted in the prolongation of certain processing stages. Shiller et al., observed the prolongation of two microstates during the IAT task: one beginning around 200 ms, with putative brain sources localized the in lingual gyrus (part of the occipital cortex specialized in letter perception), and another beginning 440 ms after stimulus onset, with putative brain sources localized in posterior cingulate and middle cingulate cortex. The latter was interpreted by the authors as increased cognitive control required during response selection, which was initiated on the subsequent microstate (Schiller et al., 2016). Similarly, microstate analysis of Stroop color-word task (Ruggeri et al., 2019) revealed that incongruent trials were characterized by slower progression through late microstates (from around 550 ms after stimulus onset), associated with response selection and execution.

From a theoretical perspective, common microstate sequence across various trial types in the extended MSIT supports the embodied cognition framework, such as the Theory of Event Coding (TEC; (Hommel, 2022; Hommel et al., 2001). TEC proposes a unified coding of perceptions and actions, rejecting the traditional distinction between stimulus-driven and response-driven conflicts (Kornblum & Lee, 1995). Instead, it postulates that the mere presentation of a stimulus requiring a response automatically triggers movement-related activity due to stimulus-response mapping. This view aligns with our findings, which demonstrate common processes in Simon (traditionally considered response-driven) and flanker (traditionally considered stimulus-driven) conflicts. Early (∼90-130 ms after stimulus onset, MS2) activation of the supplementary motor area and paracentral lobule – regions associated with movement preparation and execution – further supports this perspective (see Fig. 6 and Supplementary Table SII for putative sources for MS2). This activity coincided with the earliest activation of the visual areas and part of medial frontal cortex, overlapping with the frontal node of the ventral attention network, reflecting excitation/recruitment of exogenous attention. Importantly, this early activity was identical in all four conditions, including no-conflict, control trials (see Table 2). These findings are consistent with results reported by (Szűcs & Soltész, 2012), who showed in a numerical Stroop task, that EMG activity envelopes (in response executing muscles) began to rise at the same time in congruent, incongruent and neutral conditions, starting shortly after 200 ms post-stimulus (see Fig. 2 and S3 in Szűcs and Soltész 2012). The latency difference between our MS2 activation and EMG envelopes from (Szűcs & Soltész, 2012) agrees with the delay between cortical and muscle activation, which has been reported to range between 60-122 ms (after Van Acker et al., 2016).

### 4.2 Mapping of processing stages on the traditionally reported conflict-related ERP waves

Common finding in the ERP studies of conflict tasks were recently summarized by (Heidlmayr et al., 2020) who proposed (for a Stroop task) the neurocognitive model of the executive control course, built of sequential processes – conflict detection, interference suppression and conflict resolution – represented respectively by N2, N450 and late sustained (SP) waves. We do not see activation similar to the front-central N2 complex described in the literature (Heidlmayr et al., 2020). However, more recent studies with rigorous control conditions have shown that increase of N2 amplitude (commonly interpreted as conflict indicator) depends on trials frequencies and is absent when the incongruent and congruent conditions are equiprobable (Cheng et al., 2021; Kałamała et al., 2018). Another interpretation of the N2 function suggests its role in mismatch/novelty detection rather than in cognitive control required for conflict resolution (Tillman & Wiens, 2011). It was postulated that a pronounced positive deflection in the 200–320 ms after stimulus onset (denoted as P2 wave by authors), appears to indicate the deployment of selective attention required to resolve the flanker conflict (Kałamała et al., 2018; Korsch et al., 2016; Rey-Mermet et al., 2019a; Xu et al., 2020). The same authors (Kałamała et al., 2018) suggested that N2 might be “a frontal aspect of the P300 component” – a description that is very well fitting the N/P330 wave complex visible in our data (Supplementary Figure S3).

In our study, the earliest conflict-related effects were observed during microstate MS3, characterized by bilateral occipital negativity and fronto-central positivity, (P/N150 waves) and MS3 overlapped with the posterior negative wave, peaking ∼150 ms. It was stronger in F0 and FS than 00 and S0 trials and was also the only ERP effect that was present exclusively in trials with the flanker interference (Fig. 3A-C). A similar effect was seen in (Cheng et al., 2021) flanker study, where N1 was stronger for more confusing stimuli, built from monochromatic arrowheads as compared to color circles. In our fMRI analysis of the same task (Wojciechowski et al., 2024) we have also seen stronger activation of visual areas which differentiated F0 from S0 trials, and this difference was not explained by time-on-task effects, i.e. it was not explained by the longer response times in F0 trials. Now, thanks to ERP time resolution, we can indicate that this difference was focused within the relatively short, early (∼130-176 ms post-stimulus, MS3) stage of attentive visual processing, which was more intensive in trials with flanker interference. Increased N1 was hypothesized to reflect visual discriminative processes (Vogel & Luck, 2000) and attention allocation (Antonova et al., 2021). Indeed, MS3 was related to the brain sources not only in higher visual areas (lateral occipital cortex, cuneus and precuneus) but to increasing activity in posterior DAN nodes in bilateral parietal lobes (see Fig. 6 and Supplementary Table SII). In addition to bilateral parietal negativity, MS3 had a fronto-central positivity (P150). In the literature, this family of ERP components (P2, P2a, P150-250) was also suggested to reflect selective attention processes engaged in stimulus evaluation needed to resolve the flanker interference (Kałamała et al., 2018; Korsch et al., 2016; Rey-Mermet et al., 2019a; Xu et al., 2020). This fronto-central positivity can represent the positive pole of the active dipoles from visual areas and posterior DAN nodes, as we have not detected additional/separate anterior activity sources.

N450 is typically reported as having more negative potential on the fronto-central or centro-parietal electrodes in conflict trials. In our data, conflict trials were characterized by reduced amplitude of frontal negativity between ∼300-600 ms. There is apparently a negative wave (∼420 ms), in centro-parietal and occipital electrodes. However, microstate analysis has not detected any topographic map specific for this time window neither in our data nor in (Schiller et al., 2016) and even not in (Ruggeri et al., 2019), who analyzed Stroop data (hallmark task for N450 identification, when traditional approach to ERPs analysis is applied). In our data N/P420 wave (Supplementary Figure S3) falls on the border of two microstates (MS5 and MS6), corresponding to ERP components P/N330 and P/N500 (which could be interpreted as P3a and P3b waves). It seems that N450-like waves may be a by-product of two conflict sensitive components from the P3 family – the slower the evolution of MS5 and MS6 (P3a and P3b), the deeper the trough between them. It was already suggested that N450 wave might be specifically indexing the semantic incongruence, and in non-linguistic tasks the other waves, e.g. P3, might be conflict sensitive (Heidlmayr et al., 2020; Kałamała et al., 2018). Reduction of P3b amplitude is a common finding in cognitive control studies (Cheng et al., 2021; Kałamała et al., 2018; Korsch et al., 2016; West et al., 2005), and we saw it (in MS6) also in our data for Simon, flanker and multi-source conflict trials along with a smaller but longer P3a (MS5) for trials with flanker interference. Our microstate analysis suggests that N450 has no specific brain generator and does not reflect any separate processing-stage, thus supporting the presumption mentioned above that N450 is in fact a by-product of task-related alterations in two subsequent processes better described as P3a and P3b. Nonetheless, it may serve as a handy tool to measure interference effects on the basis of ERP difference waves, especially since its typically indicated sources include medial frontal cortex (e.g. West et al., 2012) which we show to be activated during MS5 and reaching its peak in MS6.

The last commonly reported conflict-related ERP is a late sustained potential (Chen & Melara, 2009; Donohue et al., 2016; West, 2003; West et al., 2005), which is visible in our data as a prolongation of a falling slope of P/N550 (P3b) – positive over posterior and negative over frontal areas (late window of ERP differences, Fig. 3H-K). Microstate analysis divided SP into two stages – MS7 and MS8, differentiated by the topography and brain sources. MS7 was built by dominant activity of the somatomotor network, diminishing activity of DAN, VAN and FP, and last small activations in visual cortex and precuneus. Persisted activity of the VAN node in the medial frontal cortex was the main source of MS8. It seems that the last motor decisions and commands could have been initiated during MS7, while during MS8, only their execution and observation/monitoring of outcomes took place. Both microstates had delayed centers of gravity in interference trials, however during Simon-only trials, there was a trend for an increased mean GFP of MS7 without its prolongation – as if response execution and outcome monitoring partially overlapped within this time window. In case of flanker and multi source interference, clear prolongation of MS7 (and MS8 in FS condition) indicated the spread of these processes. Stronger/longer slow potential (MS7 and MS8) seems to compensate for a less efficient/weaker processing during MS5 and MS6, for both of which we observed gradual drop of mean GFP (00>S0>F0>FS).

### 4.3 Interpretation of subsequent processing stages (microstates) in terms of large scale brain networks engagement

Consecutive microstates represented successive frames in a continuous flow of activity, starting in posterior visual areas during stimulus perception through parietal regions, to the sensory-motor and medial frontal areas during response execution and outcome monitoring (see Fig. 6).

Previous studies have demonstrated that voluntary allocation of attention, as required by our task to dissect a target digit from surrounding distractors, relies on downstream functional connectivity between DAN and visual cortex (Meehan et al., 2017). While DAN activation was observed already in our fMRI study (Wojciechowski et al., 2024), the increased time-resolution provided by EEG allowed us to discern distinct contributions of its posterior (intraparietal sulcus, IPS) and anterior (frontal eye field; FEF) nodes. During the early stage of the trial (up to ∼400 ms after stimulus onset; MS2-MS5; see Fig. 6A-D), DAN activity was confined to IPS. During MS6 (∼400-620 ms after stimulus onset; corresponding to P3b) both frontal and posterior nodes were activated (see Fig. 6E), while at the late stage of the trial (during response preparation and execution; MS7-MS8; see Fig. 6F-G) the activity persisted only in the frontal nodes.

These observations corroborate previous reports suggesting the crucial role of IPS in the selective allocation of perceptual resources (Gottlieb, 2002), while FEFs are engaged in saccade planning, execution and inhibition (Connolly et al., 2002). Findings from invasive recordings in monkeys further support the role of the IPS as target-representing area. During antisaccade tasks, the lateral intraparietal area (LIP), homologous to the human IPS, was found to represent cue location but not the direction of the actual movement (Gottlieb, 2002). The antisaccade task can be interpreted as a conflict between the reflexive tendency to gaze toward an abruptly presented stimulus and the task requirement to look away from it. Interestingly, when the intensity of this conflict was manipulated by changing stimulus luminance, activity in the IPS – particularly the right one – reflected conflict intensity, whereas the FEF did not (Anderson et al., 2008). In summary, the observed involvement of the IPS during the early stage of the trial, accompanied by strong activation in the visual network, aligns with its proposed role in stimulus-elicited processes. These processes include target identification, integration of exogenous saliency with the current set of goals and inhibition of task-irrelevant stimuli. Indeed, less effective processes at this stage (shorter activation of visual areas (MS4) was followed by longer DAN engagement (MS5, Fig. 7D) and correlated with more errors in visually demanding flanker trials (Fig. 7C). Conversely, activation of FEFs was observed during the late stage (beginning around MS6 to MS8). Notably, these microstates overlap with reaction times (average RT in no-conflict trials 535.86±61.08 vs. in multi-source conflict trials 744.33±83.56 ms). We hypothesized that the late FEF activation may reflect the execution of a saccade toward an already identified target.

The ventral attention network (VAN) was briefly activated at the beginning of the trial (MS2), remained inactive during the target selection stage, and reactivated around 338-417 ms after stimulus onset (MS6), coinciding with the cessation of activity in the posterior nodes of DAN (see Fig. 6H). Thus, the shift from DAN control towards VAN control in our task aligned with the transition from the stimulus-related to the response-related stage of the trial. This was confirmed by the correlation analysis, which indicated that RTs were prolonged when these processing stages (MS6 and MS7) were weaker (Fig.7A,B). Notably, in our data, VAN activation was primarily represented by its medial frontal node and accompanied by adjacent activation of supplementary motor area (SMA), which belongs to the somatomotor network, as well as parts of the frontoparietal network (FPN). Thus, the reaction times were more dependent on the later processing within somatomotor and attention networks. Engagement of the medial frontal cortex (MFC) is commonly reported in fMRI studies on conflict processing (Cieslik et al., 2015; Kim et al., 2010; Li et al., 2017). However, due to the low temporal resolution of this method, it remains unclear whether MFC engagement reflects conflict signaling, resolution, feedback processing, or a combination of these processes. One way to differentiate between these functions is through the timing of their occurrence. Our source analyses revealed biphasic activation of the MFC. The first activation occurred as early as 90-120 ms after trial onset (P/N110, MS2; see Fig. 6A), involving both VAN and somatomotor network (SMA). While this activation aligns with the role of “conflict signaling” (Choi et al., 2024), the MS2 did not differentiate conflict from the non-conflict control condition. Therefore, we interpret it not as conflict signaling but rather as general task signaling (Dosenbach et al., 2006), akin to the function of the ‘alerting network’ proposed by Posner and Petersen (Posner & Petersen, 1990; Xuan et al., 2016). The medial frontal cortex is one of the core regions associated with this network (Posner & Petersen, 1990; Xuan et al., 2016). The second activation was observed from 250 ms after trial onset (P/N330, MS5), continuing through response preparation and persisting until after the response (see Fig. 6D–G). This later phase suggests MFC involvement in conflict resolution and outcome monitoring.

The last two networks which were engaged during our task were FPN and DMN. The FPN, also known as a cognitive control network, is frequently reported to be activated across a wide variety of tasks requiring cognitive control (Cole & Schneider, 2007). In contrast, DMN is commonly reported to be deactivated during task execution (Fox et al., 2005), although its role as a purely task-negative network has been criticized based on evidence of task-related DMN activity (Dixon et al., 2018; Jurewicz et al., 2020; Spreng et al., 2014). While our previous fMRI study revealed widespread DMN deactivation in the prefrontal cortex (ventromedial, ventrolateral and dorsomedial), posterior cingulate cortex, as well as angular gyri, and temporal poles (see Fig. 3C in Wojciechowski et al., 2024), the ERP source analysis detected DMN activations primarily in two areas: the precuneus and part of the angular gyri (see Fig. 6C,D and Supplementary Table SII). We hypothesize that the DMN deactivations observed in fMRI may not be reflected in the ERPs and thus were not visible in our source reconstruction analysis. Previous studies have linked fMRI DMN deactivations with increase in the oscillatory alpha (Goldman et al., 2002; Laufs et al., 2003) or theta (Scheeringa et al., 2008) activity. ERP source reconstruction revealed small areas of DMN activation which were not visible in fMRI data. Despite their shared assignment to the same large-scale functional brain network the activations and deactivation reported in both analyses occur in different areas. Recent studies (Bzdok et al., 2015) have demonstrated high functional diversity of subregions within the posterior medial cortex (PMC), an area comprising both precuneus and posterior cingulate cortex. Some of this diversity is already visible in the parcellation into seven large-scale brain networks (Yeo et al., 2011), with five different networks being present in the vicinity of the posterior medial cortex (see Fig. 6). Bzdok and colleagues analyzed functional connectivity of PMC during resting state and across a wide range of tasks, identifying four subregions with distinct connectivity profiles (see Fig. 6 in (Bzdok et al., 2015). The deactivations observed in our fMRI study were localized closer to the corpus callosum and correspond most closely to the dorsal PCC, a subregion with functional connectivity to the dorsomedial prefrontal cortex and parts of angular gyrus. This subsystem aligns with the classical task-negative DMN. In contrast, the region of PMC activated in our ERP study was shown to exhibit a distinctly different pattern of connectivity including the IPS, FEF and right temporo-parietal junction. This subsystem was shown to be activated by tasks requiring attentional and executive processes, such as spatial processing, visual motion perception, the Simon task, and action inhibition (Bzdok et al., 2015).

### 4.4 Interaction of Simon and flanker interference at a stage of response processing

One of the persistent questions in the field of cognitive control concerns the processes engaged when multiple conflicts arise simultaneously. Are they resolved independently by separate, dedicated mechanisms or do they rely on the common mechanism? Our result, along with those of Schiller and Ruggeri (Ruggeri et al., 2019; Schiller et al., 2016), support a common conflict resolution/cognitive control mechanism. Specifically, the same sequence of brain activity clusters was observed for all trial types. This indicates that the same resources are shared when dealing with combined multiple conflicts, which may be ineffective and result in worse performance. Indeed, we observed a nonlinear decline in performance during combined interference trials compared to the linear model of flanker and Simon interference combination (see Fig. 2B). At the neuronal level, combined interference could evoke stronger activation (e.g. more simultaneously active neurons) or prolong the duration of activation of the same population of neurons. The measured activity may sum linearly – when there is no competition between tasks – up to the point of resource saturation, after which it becomes sublinear. Alternatively, summation could be superlinear if additional resources are recruited to handle the increased conflict demands.

In our ERP data, we observed both linear and sublinear effects. The early stage of the trial was dominated by visual attentional processing. The demand for visual attention was weaker in control and Simon trials but stronger for more complex visual stimuli building flanker conflict. The summation of brain activity at this stage was linear, indicating that the capacity of visual and attentional resources was sufficient to handle stimuli inducing multi-source conflict. Sublinear effects were observed during the late sustained potential. Specifically, SP amplitude was less increased and the dynamics of the MS6 and MS7 were less prolonged in multi-source confit trials than expected from the linear summation of Simon and flanker effects (see Fig. 5). As noted above, at the individual participant level, weaker MS6 and MS7 were associated with prolonged reaction times (in Simon and flanker conflict, respectively, Fig. 7A,B), which agrees with the interpretation of this time window as response selection/initiation stage.

The network composition of MS6 and MS7, with large contribution (or even domination in MS7) of somatomotor regions, links our result to those of (Wiesman et al., 2020) who reported superadditive increase of gamma power in superior parietal cortex in FS condition. No other nonlinearities/interactions in brain activity measures were detected in earlier studies comparing single and combined conflicts (Donohue et al., 2016; Frühholz et al., 2011a; Korsch et al., 2016; Rey-Mermet et al., 2019a; Scrivano & Kieffaber, 2022; Xie et al., 2020). Even in our fMRI study (using the same task and on the same participants) despite interaction effects in behavioral measures no such interaction was found in the BOLD signal (Wojciechowski et al., 2024). We then attributed the lack of nonlinear BOLD effect to the weak time resolution of the method. Now we see that indeed, the time window of nonlinear amplitude change was short (< 200 ms) and mostly related to dynamics of the transition from MS6 to MS7 microstate. Reasons for the reported lack of interaction between conflict types in other ERP studies could be twofold: i) low sensitivity of multivariate analyses with high number of factors (Korsch et al., 2016) and/or ii) analyses restricted to predefined latencies, and arbitrarily selected electrodes (Scrivano & Kieffaber, 2022). Note that interaction effect was in our data detected with the model analysis, and a trace of it was visible in post-hoc comparisons for Simon late-window ERP cluster (see Fig. 3H inset), but not when the window of the analysis was wider – as defined using flanker interference (F0 vs. 00, Fig. 3I inset). Taking into account the small size and short duration of this effect the analysis directly aimed to compare multi-source ERP data with the linear sum of individual conflicts might be necessary for its successful detection (e.g. FS00 vs. F0S0; see Fig. 5A).

### 4.5 Conclusions

Despite differences in task demands, Simon, flanker and multi-source conflicts shared a common sequence of processing stages, as indicated by common ERP waveforms and consistent number and order of microstates across conditions. However, the intensity and duration of these processing stages varied systematically with conflict difficulty (as measured with RTs) changing from Simon to flanker to the multi-source interference in the conflict-specific manner. Three time windows were affected by conflict presence: early (100 - 150 ms), middle (250 to over 600 ms) and late (600 to 800 ms after stimulus onset). Only the early time window was specifically modulated by trials including flanker conflict. Source analysis suggests that this stage was associated with activation in visual areas and posterior nodes of the dorsal attention network, likely due to the presence of visual distractors. When Simon and flanker conflicts were presented simultaneously, their early and middle stage neural signatures summed in a linear fashion, whereas their late-stage processing exhibited interactive effects evident in direct comparison with the additive model. Small amplitude of this effect may explain why it remained unnoticed in some earlier studies which tested it on time windows and channels predefined based on effects for other conflicts. Excellent time-resolution of ERPs allowed us to observe short scale time dynamics of large brain networks involved in our task. In line with the Theory of Event Coding we observed that mere presentation of the stimulus activated related motor programmes as indicated by early (∼90-130 ms after stimulus onset) activation of the somatomotor system (paracentral lobule and supplementary motor area). Both dorsal and ventral attention networks were involved during the trials but with different time dynamics. Dorsal attention network was most involved during the early and middle stage of the trial with activation of its posterior nodes (i.e. intraparietal sulcus) preceding activation of its frontal nodes (i.e. frontal eye fields). Ventral attention network was more engaged in the middle and late stage of the trial. Taken together, these results support a hybrid model of cognitive control, in which a common set of processing stages underlies conflict resolution, but the intensity and duration of these stages are modulated by task-specific demands.

## Acknowledgments

This work was supported by the National Science Centre, Poland; grants number UMO-2016/20/W/NZ4/00354, 2019/34/E/HS6/00257.

The authors gratefully acknowledge the help of Alicja Dobrzykowska and Olga Stefańska during EEG and MRI sessions.

The study was evaluated and approved by the ethical committee of the University of Nicolaus Copernicus in Toruń, Poland. Before taking part in the study, all subjects gave their written informed consent for participation and were screened for any contraindications to magnetic resonance imaging.

## Notes

https://openneuro.org/datasets/ds004621/versions/1.0.1

### Competing Interest Statement

The authors have declared no competing interest.

