## Supplementary materials for "Common but different: An ERP study of single- and multi-source interference processing in MSIT"

Katarzyna Paluch*, Ingrida Antonova*, Jan Nikadon, Patrycja Dzianok, Katarzyna Jurewicz, Jakub Wojciechowski, Ewa Kublik#
* equal contribution

##### Microstate cross validation procedure

Increasing numbers (3-12) of the microsate topographic maps were estimated to find the optimal one with the best fit in a cross validation procedure (see Method section). The optimal number of microstates was selected as the one for which the average increment in the percent of explained variance was lower than 0.5% (see Figure S1 below).

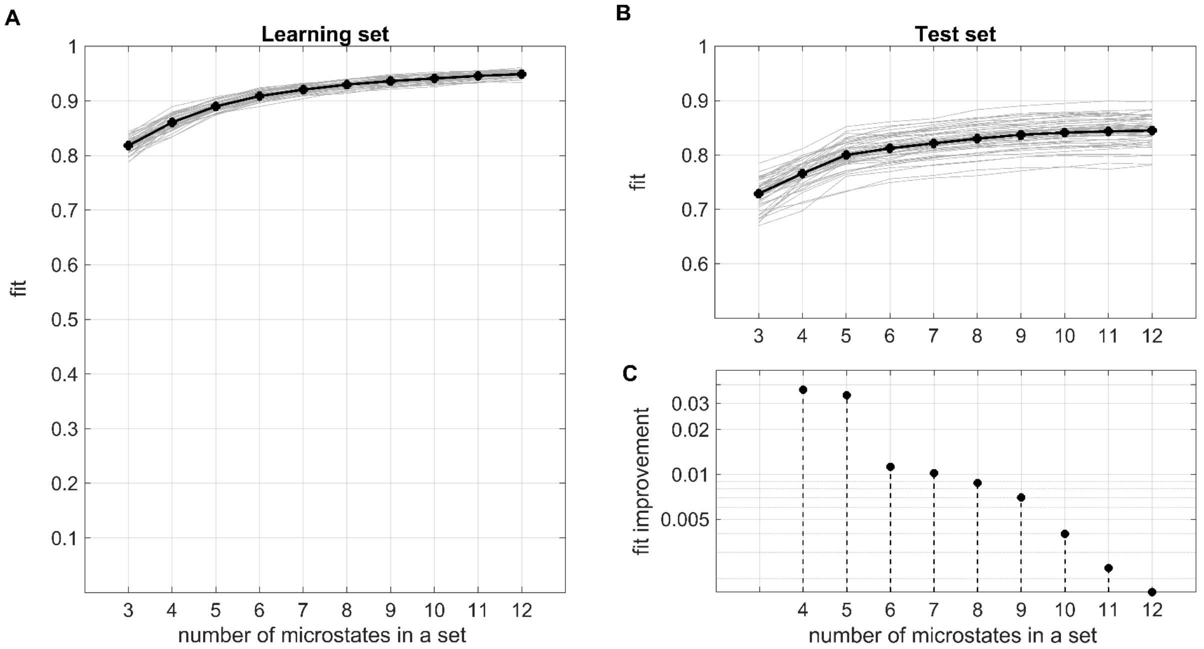


***Figure S1. The cross-validation analysis*** *indicated nine microstate clusters as the optimal set. A. Quality of fit for a learning ERP data set. B. Results indicating how well a test ERP set fits into microstates obtained in the learning set.. Bold, black lines indicate average of all iterations, thin gray lines indicate individual 250 runs. C. The average improvement of explained variance with increasing number of microstates in a set, i.e. the difference between the variance explained in a given set size minus the value for a set with n-1 size. Size of 9 microstates was chosen as the last one giving an improvement larger than 0.5%.*

**Surface maps of brain sources**

Surface maps of brain sources detected for each microstate and that of Cortical Functional Networks Atlas (Yeo et al., 2011) were prepared in BrainNet toolbox (Xia et al., 2013), with nearest neighbor voxel interpolation and BrainMesh ICBM152 smoothed template. In the case of the atlas dataset, nearest neighbor interpolation produced a surface map with discontinuities (Fig. S4 left). Figures were edited in Gimp (version 2.10.36) to enhance the saturation and fill white gaps with neighboring colors.

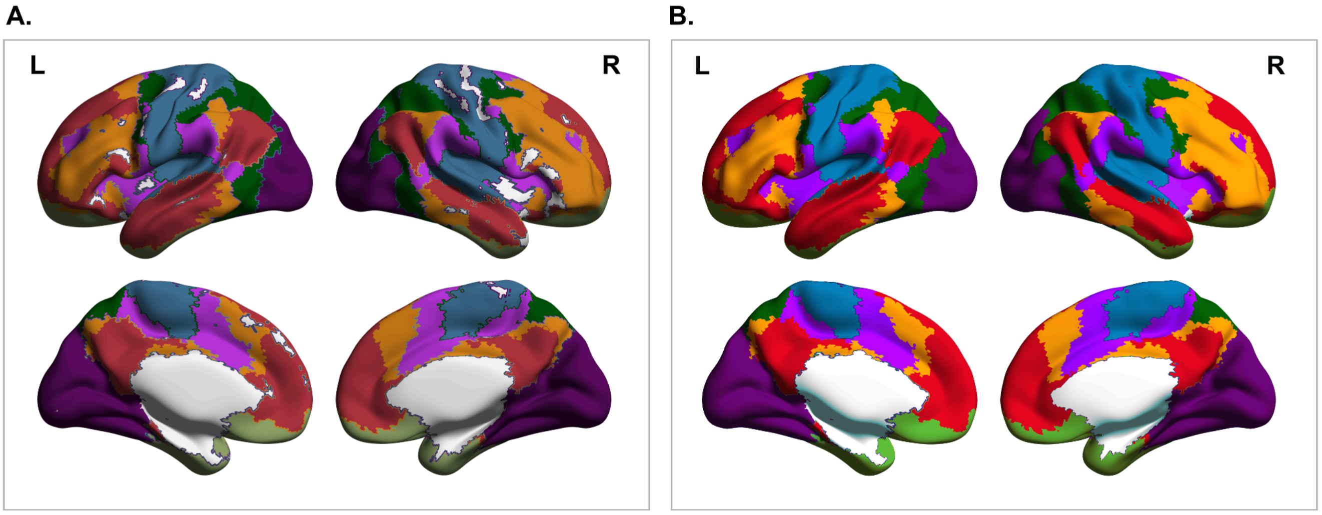


***Figure S2****. Surface map of Cortical Functional Networks – panel A. presents graphics as exported directly from BrainNet toolbox; panel B. presents a version edited for better visibility, which was used for overlay with microstate's sources in the main text Figure 6.*

#### ERP waveforms

Example grand average ERPs (from multi-source condition, FS) are shown in Figure S3. The waveforms' naming indicated in the figure (and used in the result description in the main text) is based on latency and polarity. Over time, many ERP components began to be referred to by fixed names, depending more on the context/function than on the exact time of their occurrence. We believe that attribution of these "functional names" belongs to conclusion/discussion, while in Results we describe waves by their polarity and latency features.

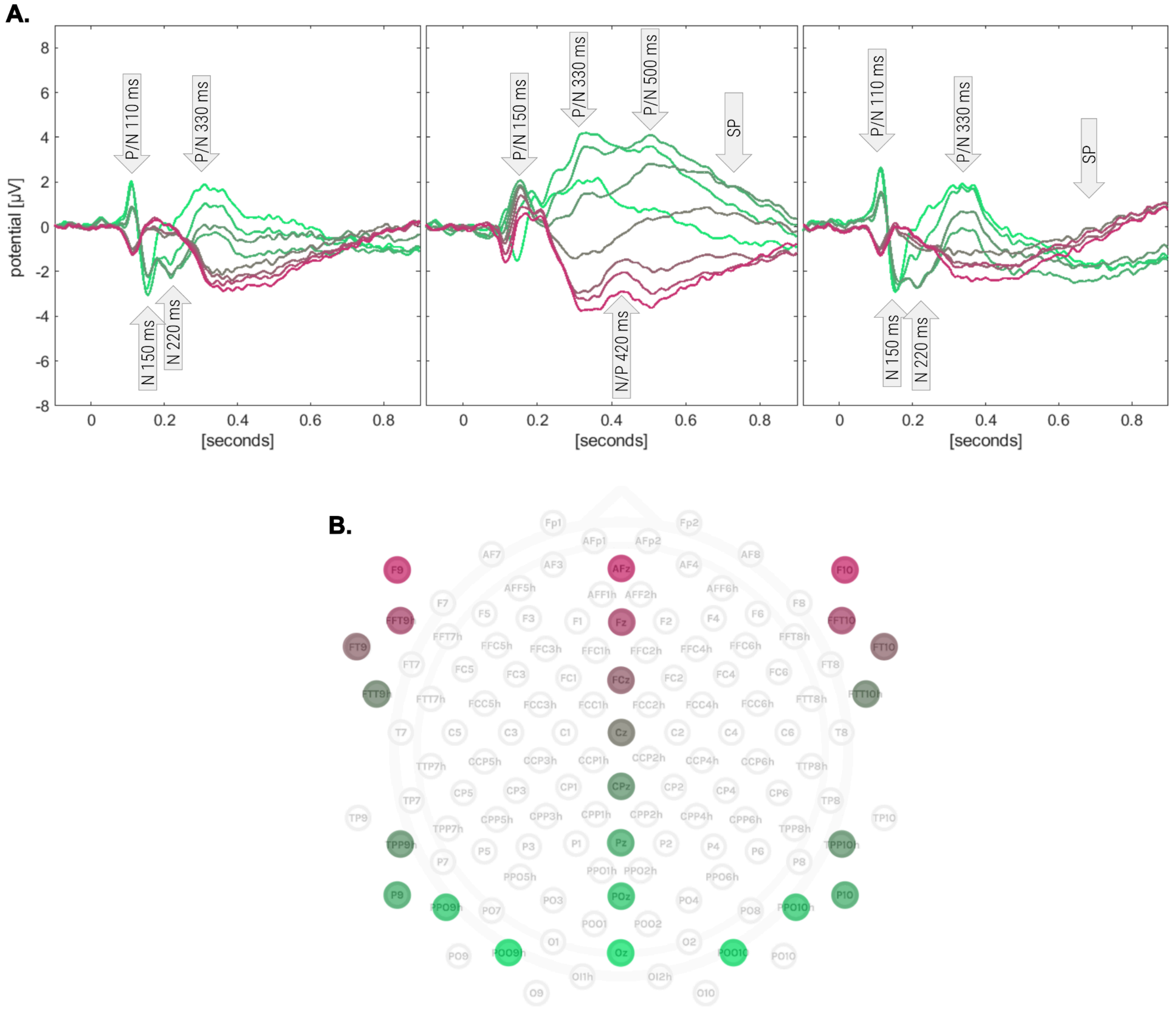


***Figure S3***. ***A.*** *Exemplary grand average ERPs (from multi-source condition, FS) highlighting individual waves seen on the ERP shape at example electrode's locations along the middle line and on the left and right side of the head (as indicated in the electrode map below). Names of individual peaks and troughs are indicated in the arrows.*
***B.*** *Schema of the electrode layout used for EEG recording, with a color legend linking particular channels to ERP waveforms in panel A. Only anterior-posterior axis is covered by a color scale because left, central and right locations are presented in corresponding left, central and right ERP plots.*

**Cluster based permutation analysis of ERP waveforms**

**Supplementary Table SI.** Results of post-hoc comparisons of cluster magnitudes (see also insets in Figure 3 and Figure 5A in the main text). Insignificant results are written in a gray font, trend-level results (0.01 < p < 0.05) in an italic font.

| | | **Post-hoc comparisons for combined posterior and anterior clusters (p-values)** | | | | | |
|---|---|---|---|---|---|---|---|
| | | Bonferroni corrected significance threshold = 0.01 | | | | | |
| | | S0-00 | F0-00 | FS-00 | F0-S0 | FS-S0 | FS-F0 |
| **Conflict versus no-conflict comparisons** | | | | | | | |
| early window N150 / P150 | S0-00 | | - | - | - | - | - |
| | F0-00 | 0.351 | | < 0.001 | < 0.001 | < 0.001 | 0.401 |
| | FS-00 | 0.269 | < 0.001 | | < 0.001 | < 0.001 | 0.923 |
| middle window N330 / P330 + N500 / P500 | S0-00 | | < 0.001 | < 0.001 | 0.115 | 0.006 | 0.076 |
| | F0-00 | *0.019* | | < 0.001 | <0.001 | < 0.001 | 0.236 |
| | FS-00 | *0.024* | < 0.001 | | 0.003 | < 0.001 | 0.104 |
| late window sustained wave over 550 ms | S0-00 | | < 0.001 | 0.105 | 0.201 | 0.405 | 0.002 |
| | F0-00 | 0.007 | | < 0.001 | < 0.001 | 0.008 | 0.463 |
| | FS-00 | *0.014* | < 0.001 | | < 0.001 | < 0.001 | 0.600 |
| | | *0.015* | < 0.001 | | *0.036* | 0.005 | *0.043* |
| **Direct F0 versus S0 comparison** | | | | | | | |
| early window | F0-S0 | 0.202 | < 0.001 | < 0.001 | | < 0.001 | 0.456 |
| middle window | | *0.023* | < 0.001 | < 0.001 | | < 0.001 | 0.493 |
| late window | | 0.008 | 0.006 | *0.034* | | 0.523 | 0.542 |
| **Interaction (FS00 vs. F0S0)** | | Bonferroni corrected significance threshold = 0.0083 | | | | | |
| early window | FS00-F0S0 | - | - | - | - | - | - |
| middle window | | - | - | - | - | - | - |
| late window | | 0.005 | < 0.001 | 0.179 | 0.113 | 0.565 | 0.005 |

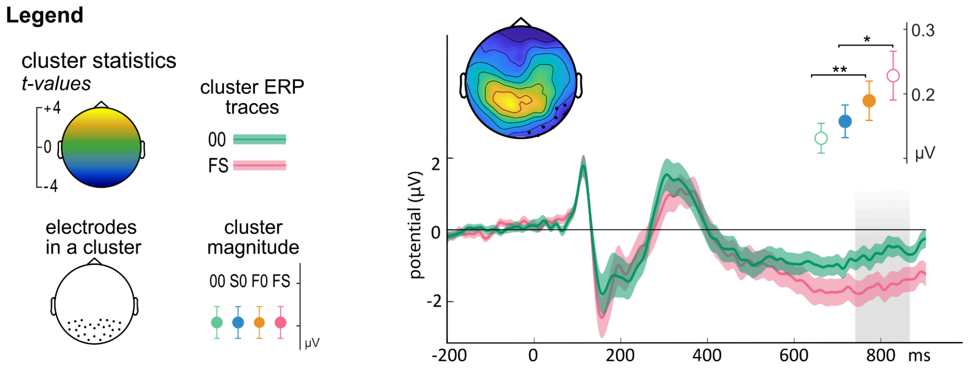


***Figure S4.*** *Average ERP waveforms, difference map and results of post-hoc test for a second activity cluster detected in late window FS versus 00 comparison (right occipito-parietal negativity: 737-862 ms, 8 electrodes, p = 0.028)). Green waveform - 00, red waveform - FS; gray shading - time window of significant difference defining cluster duration. Small dot-chart presents mean absolute amplitudes of a cluster window in all four MSIT conditions (green 00; blue S0; yellow F0, red FS). Asterisk mark significance from post-hoc paired comparisons of cluster magnitude: F0 vs 00, p = 0.001 (**); S0 vs FS p = 0.036(*). The empty dots mark cluster-defining comparison, which was not included in post-hoc tests. Significance threshold after Bonferoni correction for 5 comparisons: p = 0.01.*

**Results of source analysis**

Detailed information about detected activity clusters is presented in the Supplementary Tables SII, below. The activity cluster characteristics include: MNI Coordinates; Cluster size (i.e. number of voxels); anatomical regions (according to AAL atlas) and a % of cluster they occupied, and a % of a whole region that was active within the cluster; statistical significance of a cluster with (FWE and FDR corrections for multiple comparisons) and without correction for multiple comparison, and a peak t-test value.

**Common but different: An ERP study of single- and multi-source interference processing in MSIT**
**Supplementary Table SII**

| Coordinates MNI x | y | z | Cluster size | Anatomical region (AAL atlas) | % of cluster | % or region | Cluster pFWE | Cluster pFDR | Cluster p (uncor) | Peak T |
|---|---|---|---|---|---|---|---|---|---|---|
| **MS2** | | | | | | | | | | |
| | | | | Precuneus_L | 7,8 | 48,4 | | | | |
| | | | | Calcarine_L | 7,7 | 74,3 | | | | |
| | | | | Calcarine_R | 7,1 | 82,6 | | | | |
| | | | | Cuneus_L | 6,6 | 95 | | | | |
| | | | | Cuneus_R | 6,5 | 99,4 | | | | |
| | | | | Occipital_Mid_R | 6,1 | 63,4 | | | | |
| | | | | Occipital_Sup_L | 6 | 96,4 | | | | |
| **20** | **-84** | **34** | **21800** | **Occipital_Sup_R** | **5,7** | **87,8** | **< 0.001** | **< 0.001** | **< 0.001** | **8,06** |
| | | | | Precuneus_R | 5,6 | 37,4 | | | | |
| | | | | Occipital_Mid_L | 4,4 | 29,4 | | | | |
| | | | | Parietal_Sup_L | 4,1 | 43,7 | | | | |
| | | | | Lingual_R | 4,1 | 38,9 | | | | |
| | | | | LIngual_L | 3,9 | 40,4 | | | | |
| | | | | Parietal_Sup_R | 1,2 | 12 | | | | |
| | | | | **sum** | **76,8** | | | | | |
| | | | | OUTSIDE | 20,9 | | | | | |
| **6** | **14** | **34** | **3934** | **Cingulate_Mid_R** | **20,1** | **35,9** | **< 0.001** | **< 0.001** | **< 0.001** | **4,44** |
| | | | | Cingulate_Mid_L | 15,8 | 32,1 | | | | |
| | | | | Supp_Motor_Area_L | 15,8 | 29 | | | | |
| | | | | Supp_Motor_Area_R | 11,6 | 19,3 | | | | |
| | | | | Paracentral_Lobule_L | 6,2 | 18,2 | | | | |
| | | | | ACC_sup_R | 1,8 | 13,1 | | | | |
| | | | | Frontal_Sup_2_L | 1 | 0,8 | | | | |
| | | | | **sum** | **72,3** | | | | | |
| | | | | OUTSIDE | 25,6 | | | | | |

| Coordinates MNI x | y | z | Cluster size | Anatomical region (AAL atlas) | % of cluster | % or region | Cluster pFWE | Cluster pFDR | Cluster p (uncor) | Peak T |
|---|---|---|---|---|---|---|---|---|---|---|
| **MS3** | | | | | | | | | | |
| | | | | Occipital_Mid_L | 8,7 | 69,3 | | | | |
| | | | | Precuneus_R | 5,2 | 41,5 | | | | |
| **32** | **-68** | **28** | **25956** | **Occipital_Mid_R** | **5,1** | **63,2** | **< 0.001** | **< 0.001** | **< 0.001** | **5,74** |
| | | | | Cerebelum_6_L | 4,5 | 68,4 | | | | |
| | | | | Cuneus_R | 4,4 | 79,9 | | | | |
| | | | | Lingual_L | 4,2 | 52,7 | | | | |
| | | | | Occipital_Sup_R | 4,1 | 75,8 | | | | |
| | | | | Calcarine_R | 4 | 56 | | | | |
| | | | | Calcarine_L | 3,8 | 43,5 | | | | |
| | | | | Cerebelum_Crus1_L | 3,6 | 36 | | | | |
| | | | | Fusiform_L | 3,5 | 39,3 | | | | |
| | | | | Occipital_Sup_L | 3,3 | 61,8 | | | | |
| | | | | Precuneus_L | 3,1 | 23 | | | | |
| | | | | Cuneus_L | 3 | 51,7 | | | | |
| | | | | Lingual_R | 2,7 | 30 | | | | |
| | | | | Occipital_Inf_L | 1,8 | 49,1 | | | | |
| | | | | Parietal_Sup_R | 1,6 | 18,7 | | | | |
| | | | | Parietal_Sup_L | 1,6 | 20,1 | | | | |
| | | | | Angular_R | 1,5 | 22 | | | | |
| | | | | Fusiform_R | 1,4 | 14,5 | | | | |
| | | | | Angular_L | 1,3 | 28,4 | | | | |
| | | | | Temporal_Mid_L | 1,3 | 6,6 | | | | |
| | | | | Temporal_Mid_R | 1,2 | 6,9 | | | | |
| | | | | **sum** | **74,9** | | | | | |
| | | | | OUTSIDE | 20% | | | | | |

| Coordinates MNI | | | Cluster size | Anatomical region (AAL atlas) | % of cluster | % or region | Cluster pFWE | Cluster pFDR | Cluster p (uncor) | Peak T |
|---|---|---|---|---|---|---|---|---|---|---|
| x | y | z | | | | | | | | |
| **MS4** | | | | | | | | | | |
| | | | | Occipital_Mid_L | 6,8 | 68,9 | | | | |
| | | | | Precuneus_R | 6,2 | 63,2 | | | | |
| **36** | **-70** | **28** | **33222** | **Occipital_Mid_R** | **5,1** | **81,3** | **< 0.001** | **< 0.001** | **< 0.001** | **7,84** |
| | | | | Temporal_Mid_R | 5,1 | 38,7 | | | | |
| | | | | Angular_R | 4,5 | 85,2 | | | | |
| | | | | Calcarine_R | 4,4 | 78,7 | | | | |
| | | | | Temporal_Mid_L | 3,8 | 25,5 | | | | |
| | | | | Fusiform_R | 3,7 | 48,7 | | | | |
| | | | | Temporal_Inf_R | 3,5 | 32,3 | | | | |
| | | | | Cuneus_R | 3,4 | 79,6 | | | | |
| | | | | Occipital_Sup_R | 3,4 | 79 | | | | |
| | | | | Lingual_R | 2,8 | 40,8 | | | | |
| | | | | Parietal_Sup_R | 2,6 | 39,1 | | | | |
| | | | | Precuneus_L | 2,5 | 23,8 | | | | |
| | | | | Temporal_Inf_L | 2,1 | 22,1 | | | | |
| | | | | Fusiform_L | 2 | 28,1 | | | | |
| | | | | Cuneus_L | 1,6 | 34,5 | | | | |
| | | | | Occipital_Inf_L | 1,5 | 51,3 | | | | |
| | | | | Calcarine_L | 1,3 | 19,7 | | | | |
| | | | | Cerebelum_6_R | 1,1 | 21,1 | | | | |
| | | | | Parietal_Sup_L | 1 | 16 | | | | |
| | | | | Cerebelum_Crus1_R | 1 | 12,1 | | | | |
| | | | | **sum** | **69,4** | | | | | |
| | | | | (24%) OUTSIDE | 24,2 | | | | | |

| Coordinates MNI | | | Cluster size | Anatomical region (AAL atlas) | % of cluster | % or region | Cluster pFWE | Cluster pFDR | Cluster p (uncor) | Peak T |
|---|---|---|---|---|---|---|---|---|---|---|
| x | y | z | | | | | | | | |
| **MS5** | | | | | | | | | | |
| | | | | Precuneus_R | 9 | 66,9 | | | | |
| | | | | Occipital_Mid_L | 6,7 | 50,2 | | | | |
| | | | | Precuneus_L | 6,1 | 42,4 | | | | |
| | | | | Parietal_Sup_L | 5,9 | 69,4 | | | | |
| | | | | Occipital_Mid_R | 5,1 | 59 | | | | |
| | | | | Parietal_Sup_R | 5,1 | 55,5 | | | | |
| | | | | Parietal_Inf_L | 4,9 | 49 | | | | |
| | | | | Cuneus_R | 4,1 | 70 | | | | |
| **30** | **-72** | **42** | **24360** | **Occipital_Sup_R** | **4,1** | **70** | **< 0.001** | **< 0.001** | **< 0.001** | **6,23** |
| | | | | Cuneus_L | 3,8 | 61,5 | | | | |
| | | | | Angular_L | 3,7 | 76,2 | | | | |
| | | | | Temporal_Mid_R | 2,8 | 15,6 | | | | |
| | | | | Angular_R | 2,7 | 37,8 | | | | |
| | | | | Occipital_Sup_L | 2,5 | 44,6 | | | | |
| | | | | Temporal_Mid_L | 2,3 | 11,5 | | | | |
| | | | | Paracentral_Lobule_R | 1,7 | 48,7 | | | | |
| | | | | Postcentral_L | 1,6 | 10,02 | | | | |
| | | | | Cingulate_Mid_L | 1,6 | 19,6 | | | | |
| | | | | Supp_Motor_Area_L | 1,4 | 15,8 | | | | |
| | | | | Frontal_Sup_2_L | 1,1 | 5,5 | | | | |
| | | | | Postcentral_R | 1 | 6,1 | | | | |
| | | | | **sum** | 77,2 | | | | | |
| | | | | OUTSIDE | 19,8 | | | | | |

| Coordinates MNI x | y | z | Cluster size | Anatomical region (AAL atlas) | % of cluster | % or region | Cluster pFWE | Cluster pFDR | Cluster p (uncor) | Peak T |
|---|---|---|---|---|---|---|---|---|---|---|
| **MS6** | | | | | | | | | | |
| | | | | Precuneus_L | 8,5 | 73,2 | | | | |
| | | | | Supp_Motor_Area_R | 6,8 | 86,9 | | | | |
| | | | | Parietal_Sup_L | 6,3 | 91,8 | | | | |
| | | | | Supp_Motor_Area_L | 5,5 | 78 | | | | |
| | | | | Postcentral_L | 4,4 | 34 | | | | |
| | | | | Cuneus_L | 4,3 | 85,3 | | | | |
| | | | | Parietal_Inf_L | 3,9 | 48,4 | | | | |
| | | | | Cingulate_Mid_L | 3,6 | 56 | | | | |
| | | | | Occipital_Mid_L | 3,5 | 32,7 | | | | |
| | | | | Cingulate_Mid_R | 3,5 | 47,6 | | | | |
| | | | | Precuneus_R | 3,3 | 30,1 | | | | |
| **-18** | **-76** | **28** | **30265** | **Occipital_Sup_L** | **3,2** | **71,9** | **< 0.001** | **< 0.001** | **< 0.001** | **6,68** |
| | | | | Precentral_L | 3,2 | 27,5 | | | | |
| | | | | Calcarian_L | 3,1 | 41,7 | | | | |
| | | | | Paracentral_Lobule_L | 2,2 | 49,2 | | | | |
| | | | | Occipital_Mid_R | 2 | 28,2 | | | | |
| | | | | Frontal_Sup_2_R | 1,9 | 11,3 | | | | |
| | | | | Cuneus_R | 1,8 | 39 | | | | |
| | | | | Angular_L | 1,6 | 41,4 | | | | |
| | | | | Occipital_Sup_R | 1,3 | 28,3 | | | | |
| | | | | Calcarine_R | 1,3 | 21 | | | | |
| | | | | Frontal_Sup_2_L | 1,2 | 7,7 | | | | |
| | | | | **sum** | **76,4** | | | | | |
| | | | | OUTSIDE | 20,6 | | | | | |

| Coordinates MNI x | y | z | Cluster size | Anatomical region (AAL atlas) | % of cluster | % or region | Cluster pFWE | Cluster pFDR | Cluster p (uncor) | Peak T |
|---|---|---|---|---|---|---|---|---|---|---|
| **MS7** | | | | | | | | | | |
| | | | | Supp_Motor_Area_L | 10,8 | 79,8 | | | | |
| **-22** | **-18** | **64** | **15863** | **Precentral_L** | **7,7** | **34,8** | **< 0.001** | **< 0.001** | **< 0.001** | **5,01** |
| | | | | Supp_Motor_Area_R | 7,6 | 50,6 | | | | |
| | | | | Cingulate_Mid_R | 7 | 50,2 | | | | |
| | | | | Postcentral_L | 6,2 | 25,2 | | | | |
| | | | | Frontal_Sup_2_L | 5,7 | 18,5 | | | | |
| | | | | Frontal_Sup_2_R | 3,7 | 11,5 | | | | |
| | | | | Cingulate_Mid_L | 2,8 | 22,6 | | | | |
| | | | | Parietal_Sup_L | 2,6 | 20 | | | | |
| | | | | Parietal_Inf_L | 1,8 | 11,6 | | | | |
| | | | | Frontal_Mid_2_L | 1,4 | 4,8 | | | | |
| | | | | Paracentral_Lobule_L | 1,2 | 13,8 | | | | |
| | | | | Frontal_Mid_2_R | 1,1 | 3,5 | | | | |
| | | | | Caudate_R | 1 | 18,3 | | | | |
| | | | | **sum** | **60,6** | | | | | |
| | | | | OUTSIDE | 37,4 | | | | | |
| | | | | Calcarian_R | 54,6 | 21,6 | | | | |
| **18** | **-76** | **18** | **734** | **Cuneus_R** | **25,8** | **13,3** | **0.055** | **0.008** | **0.008** | **4,05** |
| | | | | Occipital_Sup_R | 12,7 | 6,6 | | | | |
| | | | | Precuneus_R | 1,1 | 0,3 | | | | |
| | | | | **sum** | **94,2** | | | | | |
| | | | | OUTSIDE | 5 | | | | | |

| Coordinates MNI x | y | z | Cluster size | Anatomical region (AAL atlas) | % of cluster | % or region | Cluster pFWE | Cluster pFDR | Cluster p (uncor) | Peak T |
|---|---|---|---|---|---|---|---|---|---|---|
| **MS8** | | | | | | | | | | |
| **-2** | **16** | **46** | **10787** | **Supp_Motor_Area_L** | **10,7** | **53,8** | **< 0.001** | **< 0.001** | **< 0.001** | **4,88** |
| | | | | Cingulate_Mid_R | 9,9 | 48,5 | | | | |
| | | | | Supp_Motor_Area_R | 8,5 | 38,6 | | | | |
| | | | | Frontal_Sup_2_L | 6,9 | 15,3 | | | | |
| | | | | Precentral_L | 5,5 | 16,8 | | | | |
| | | | | Cingulate_Mid_L | 4,5 | 25,2 | | | | |
| | | | | Frontal_Sup_2_R | 3,2 | 6,8 | | | | |
| | | | | Frontal_Mid_2_L | 2 | 4,7 | | | | |
| | | | | Frontal_Sup_Medial_L | 1,6 | 5,7 | | | | |
| | | | | Caudate_R | 1,5 | 19,1 | | | | |
| | | | | **sum** | **54,3** | | | | | |
| | | | | OUTSIDE | 43,9 | | | | | |
